# Quantitative Control of Noise in Mammalian Gene Expression by Dynamic Histone Regulations

**DOI:** 10.1101/2020.12.20.423693

**Authors:** Deng Tan, Rui Chen, Yuejian Mo, Wei Xu, Xibin Lu, Huiyu He, Shu Gu, Fan Jiang, Weimin Fan, Yilin Wang, Xi Chen, Wei Huang

**Affiliations:** Department of Biology, Southern University of Science and Technology, Shenzhen, Guangdong, 518055, China; Core Research Facilities, Southern University of Science and Technology, Shenzhen, Guangdong, 518055, China; Department of Biomedical Engineering, Southern University of Science and Technology, Shenzhen, Guangdong, 518055, China

## Abstract

Fluctuation (‘noise’) in gene expression is critical for mammalian cellular processes. Numerous mechanisms contribute to its origins, yet large noises induced by single transcriptional activator species remain to be experimentally understood. Here, we combined the dynamic regulation of transcriptional activator binding, histone regulator inhibitors, and single-cell quantification of chromatin accessibility, mRNA, and protein to probe putative mechanisms. Using a light-induced expression system, we show that the transcriptional activator forms a positive feedback loop with histone acetyltransferases CBP/p300. It generates epigenetic bistability in H3K27ac, which contributes to large noise. Disable of the positive feedback loop by CBP/p300 and HDAC4/5 inhibitors also reduces heterogeneity in endogenous genes, suggesting a universal mechanism. We showed that the noise was reduced by pulse-wide modulation of transcriptional activator binding due to alternating the system between high and low monostable states. Our findings could provide a mechanism-based approach to modulate noise in synthetic and endogenous gene expressions.

## Introduction

Isogenic cell in a homogenous environment exhibit significant variations in gene expression. A single cell also shows similar gene expression fluctuation over time. The phenomenon, often called gene expression noise, was initially demonstrated by computational modeling of stochastic biochemical reactions with finite biomolecules (McAdams and Arkin, 1997). Elowitz *et al*. started experimental discoveries of its prevalence (Elowitz et al., 2002). Many general mechanisms have been identified to contribute to gene expression noise in mammalian cell, including partition at cell division, transcriptional bursting, epigenetic modifications, 3D chromosome structure, *etc*. (Huh and Paulsson, 2011; Nicolas et al., 2018; Rodriguez et al., 2019; Singer et al., 2014; Suter et al., 2011). However, the mechanistic origin of large noise caused by single transcriptional activator in mammalian cell remains to be identified experimentally. On the other hand, various biological processes such as cell fate control in proliferation, differentiation, and cell death, rely on precise control of gene expression (Balázsi et al., 2011). There are cases of cells and animals suppressing or elevating noises to benefit their performance objectives (Chang et al., 2008; Hansen et al., 2018; Li et al., 2017; Sosnik et al., 2016). Synthetic inducible gene expression systems have been developed to perform designed perturbations of biological processes in cells and animals, including the original Tet-Off (Gossen and Bujard, 1992), Tet-On (Gossen et al., 1995), other chemicals (Khalil et al., 2012), light-induced (Wang et al., 2012), and CRISPR-derived ones (Nihongaki et al., 2017; Shao et al., 2018). Together with other toolboxes developed in synthetic biology, we can further understand the complex biological processes with quantitative and dynamic modulation of critical gene products in mammals. They also enable studies of the basic process of transcription regulations with such dynamic and quantitative modulations, not readily possible with endogenous gene regulations.

An inducible expression system usually consists of an engineered transcriptional activator and an engineered promoter with a minimal promoter (TATA box) and multiple binding sites for the transcriptional activators. A transcriptional activator was usually constructed with a DNA binding domain and an activation domain of endogenous transcriptional activator such as nuclear factor kappa-B (NF κ-B) p65 subunit (p65AD) or VP16 (VP16AD) from herpes simplex virus type I. Upon bound to the promoter, the transcriptional activator recruits co-activators, mediator complex and transcriptional preinitiation complex (PIC) to initiate transcription. Such simple gene expression systems often exhibit significant larger noises with unclear mechanisms. It also limited the potential applications in studies of cell fate control. Several engineering approaches have been reported to suppress noises in doxycycline- (dox-) and light-induced expressions in mammalian cell. Two engineered circuits constructed with dox-induced tetR-based negative feedback loop (Nevozhay et al., 2013) and light-induced tetR-LOV negative feedback loop (Guinn and Balázsi, 2019) were demonstrated to constitutively reduce noises in HEK293 cell. Two fundamental properties of these hardwired circuits limited their potential applications: fixed noise levels and cell-type sensitive CMV promoter.

The present study is to identify potential mechanisms of large noises generated by single transcriptional activator utilizing a lightON inducible gene expression (Wang et al., 2012), and develop strategies to independently control noise and mean expression. We generated stably transfected human ovarian cancer (HeLa), and mouse embryonal carcinoma (F9) cell clones with this lightON expression system incorporated a p65AD and implemented dynamic and quantitative light inductions. We used flow cytometry to discover that pulse-width modulation (PWM) illumination with a period of 400 min or longer reduces gene expression noise to a basal level. We identified that histone acetylation is the key to the large noises in the amplitude modulation (AM) illumination. We hypothesize that light-induced transcriptional activator binding to the promoter in a stochastically opened chromatin, recruiting the CBP/p300 co-activators. The CBP/p300 not only facilitates the recruitment of PIC and initiate transcription, but also acetylate histones in the vicinity. The acetylated histones help keep the chromatin at an active/open state. At intermediate AM illuminations, this positive feedback loop could generate bistability in histone acetylation and transcriptional activity. The PWM induction reduces noise by alternating the cell between “high” and “low” monostable states. We used A485, a specific inhibitor for CBP/p300 HAT activities (Lasko et al., 2017), H3K27ac ChIP-seq analysis, and single-cell transposase-accessible chromatin using sequencing (scATAC-seq) analysis (Chen et al., 2018), as well as live-cell single mRNA imaging analysis to validate the hypothesis. Furthermore, we combined CBP/p300 inhibitor (A485) and HDAC4 inhibitor (LMK-235) to disrupt potential positive feedback loop involving CBP/p300 in mouse embryonic stem (mES) cell, HeLa and F9 cell, and observed the reduction of gene expression heterogeneity in genes with high noises, suggesting that it could be widely occurred.

## Results

### Modulation of gene expression noise with periodic induction in a mammalian light-inducible gene expression system

To facilitate the dynamic induction of gene expression, we generated stable clones of the lightON expression system (Wang et al., 2012) (Figure 1-figure supplement 1A) into HeLa cell using PiggyBac transposon (Lu and Huang, 2014). It consists of a synthetic transcriptional activator GAVPO, and mRuby3 (Bajar et al., 2016) reporter driven by a synthetic 5xUAS promoter consisted of 5 GAL4 binding elements and a TATA box (plasmid B1 in Figure 1-figure supplement 1B). To assess the contribution of GAVPO expression variation to the mRuby noise, the GAVPO is driven by a noise reduction circuit of CMV-tetO2-tetR-GFP-T2A-GAVPO (Nevozhay et al., 2013) to facilitate an indirect single-cell measurement of GAVPO expressions (plasmid A1 in Figure S1B). The engineered HeLa cell clone is named as HeLa-AB1. With a constant light illumination (amplitude modulation, AM) gradually increased from 0 to 100 μW/cm^2^, the population averages of mRuby expressions increase gradually (blue line in Figure 1A). The cell population exhibits a bell-shaped noise profile, quantified with coefficients of variance (CV), corresponding to light intensities (blue line in Figure 1B) and mean mRuby expressions (Figure 1C). The largest noise level (CV ∼2.5) is induced with the intermediate light intensities (∼10 μW/cm^2^). The histograms indicate that there are possibly “low” and “high” states of mRuby expressions (Solid lines in Figure 1D). The single cell expression distribution shows that more and more cells transit from the “low” to the “high” state when light intensity increases from 10 to 45 μW/cm^2^. The peaks of the low and high states don’t appear to change much. Similar phenomenon also exhibits in the dox-inducible (Tet-On) system (solid line in Figure 2-figure supplement 2B). To assess the contributions of the GAVPO noise to mRuby distribution dispersion, we used the mRuby-GFP flow cytometry data at 25 μW/cm^2^ (Figure 1E) to estimate the population average dose-response curve of mRuby versus GFP-GAVPO, which is rather shallow and not yet plateaued (Figure 1F). With limited noise of GFP-GAVPO, this analysis demonstrates that the wide distribution of mRuby is not caused by the sharp GAVPO-mRuby dose-response curve, a hypothesis supported by the recent study of light-induce gene expression noise in yeast (Benzinger and Khammash, 2018). This type of expression distribution could help multipotential stem cell explore their full differentiation potentials (Chang et al., 2008), but strategies to reduce such gene expression variations are necessary to study precise cell fate control.

**Figure 1.**
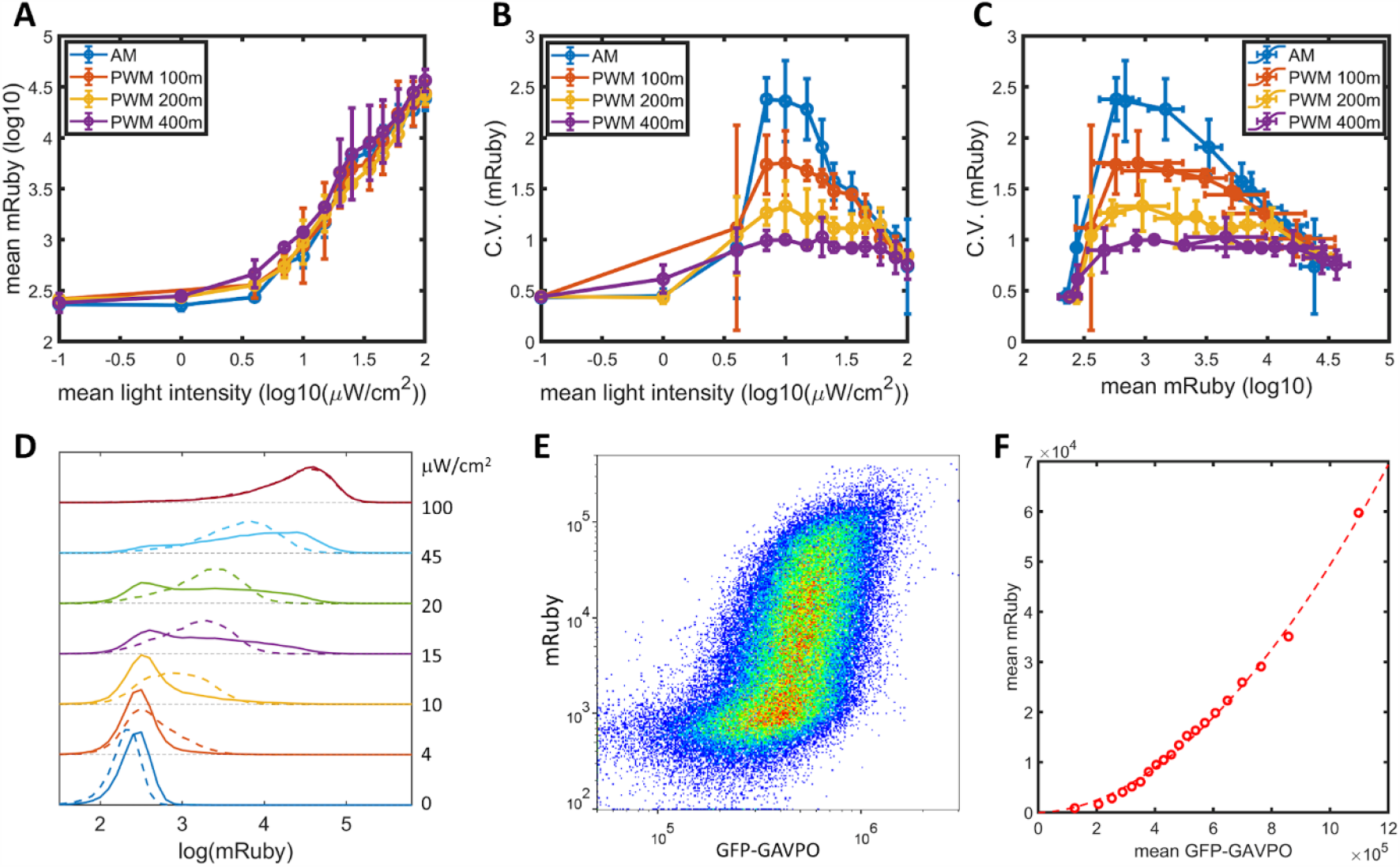
AM and PWM light induction of gene expressions and noises of lightON-mRuby expression in HeLa-AB1 cell. (**A**) The response curves of mean mRuby expressions over mean light intensities for AM (blue), and PWM with a period of 100 m (red), 200 m (yellow), and 400 m (purple). (**B-C**) The correlation curves of CV of mRuby expression versus mean light intensities (B), and mean mRuby expressions (C) for AM (blue), and PWM with a period of 100 m (red), 200 m (yellow), and 400 m (purple). The error bars represent standard deviations from 2 to 4 independent experiments. (**D**) Histogram analysis of AM modulated expression (solid lines) and PWM modulated expression (dashed lines, with a period of 400 m) with increasing mean light intensities. Specific mean light intensities are marked to the right to each histograms. (**E**) The density plot of mRuby vs GFP-GAVPO under AM with 25 μW/cm^2^. (**F**) The cells from E are divided into 20 equal populations in the ascending order of GFP intensities, and the mean mRuby versus mean GFP-GAVPO was plotted in linear scale (open circle) and fitted to a Hill function (dashed line) with hill coefficient of 1.87.

**Figure 2.**
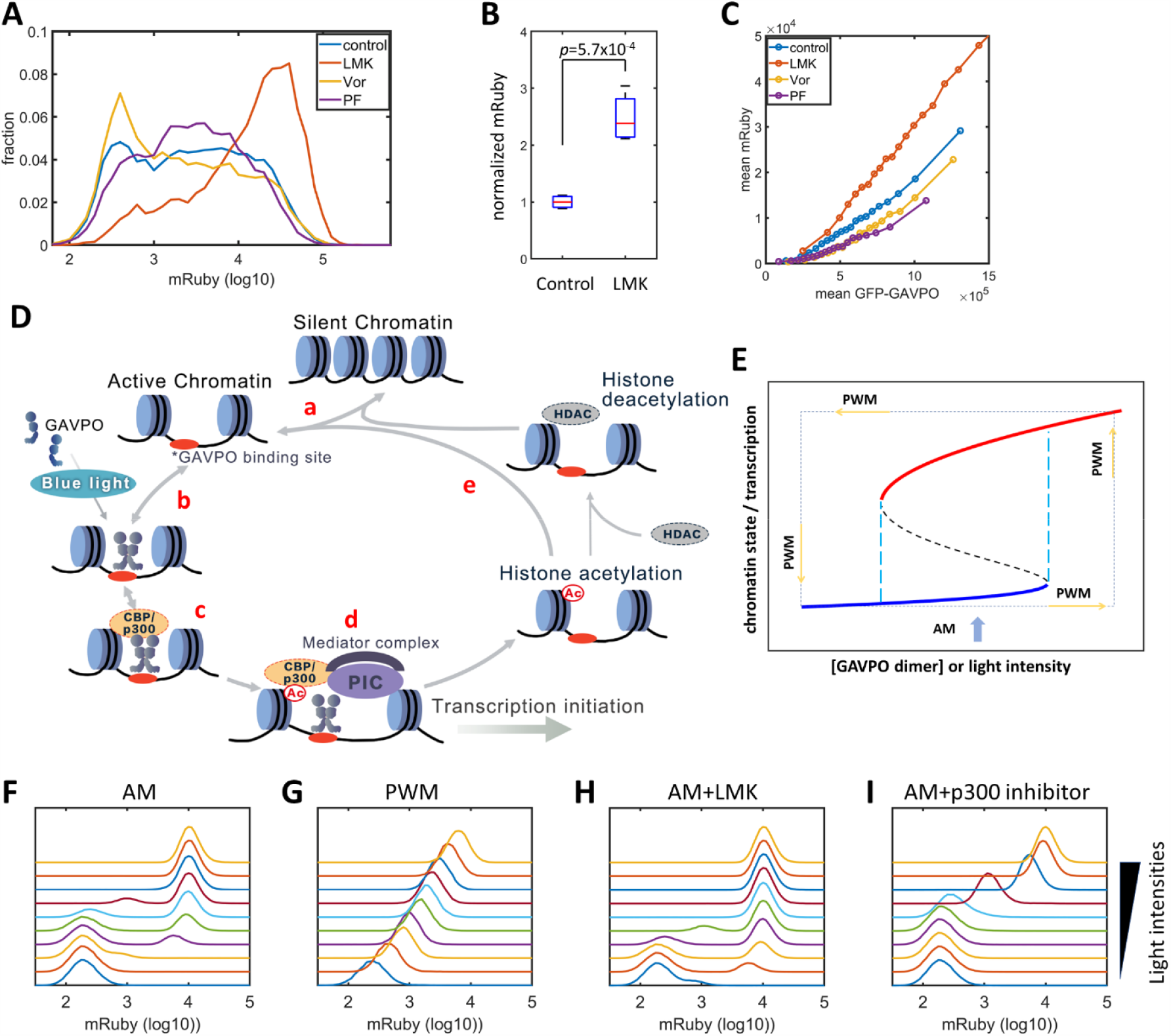
Model of light-induced gene expressions and epigenetic bistability. (**A**) Histograms of mRuby for HeLa-AB1 cells under AM with 25 μW/cm^2^, and treated with inhibitors for various epigenetic regulators. LMK, LMK-235, a selective HDAC4/5 inhibitor. Vor, Vorinostat (SAHA), a broad-spectrum inhibitor for HDACs. PF, PF-06726304, a selective inhibitor for the EZH2 component of the Polycomb Repressive Complex 2 (PRC2). The concentrations for the three inhibitors are 0.333 μM. (**B**) Box plot and unpaired *t*-test for normalized mRuby of HeLa-AB1 with control and LMK-235 treatment. Data represent 4 independent experiments. The boxes show the lower and upper quartiles, the whiskers show the minimum and maximal values excluding outliers; the line inside the box indicates the median. (**C**) The response curves of population average mRuby expressions versus population mean GFP-GAVPO expression under inhibitors to various epigenetic modifiers. (**D**) Proposed mechanism for transcription and epigenetic regulation events for lightON expressions. HDAC, HDAC4/5, and possibly other HDACs. PIC, transcriptional preinitiation complex. Ac, histone acetylation. (**E**) Schematic view of positive feedback loop-induced bistability to illustrate the AM induction of large noise and PWM reduction of noise. The black dashed lines represent the boundary that separates the high and low states. The cyan dashed lines represent the thresholds between low monostable state (blue) and bistable states (blue and red), and between bistable states (blue and red) and high monostable state (red). Light blue arrows indicate AM with intermediate intensities, that located at the bistability region. The dashed rectangle represents the cycle of PWM alternating cells between high monostable and low monostable states. (**F-I**) Simulated histograms of mRuby expressions for increasing mean light intensities for AM modulation alone (F), PWM modulation with 400 min (G), AM modulation with LMK-235(H), AM modulation with inhibitor for HAT activities of CBP/p300 (I). Each sample contains 10,000-50,000 cells.

The noise might be related to “off” and “on” epigenetic states at the 5xUAS promoter. The concept of epigenetic bistability was theorized by Sneppen *et al*. (Dodd et al., 2007; Sneppen and Ringrose, 2019). Still, experimental studies mostly contribute it to long term epigenetic memory (Hathaway et al., 2012; Singer et al., 2014), not to fast gene expression fluctuations. Bintu *et al*. demonstrated that the kinetics of different histone modifiers varies from hours to days in CHO cell (Bintu et al., 2016). We hypothesized that by periodical modulating the light intensity with proper time scales, the transcription dynamics might be entangled with histone modification dynamics to modulate gene expression noise. To explore this possibility, we use pulse-width modulation (PWM) regiment with light switching off and on (between 0 and 100 μW/cm^2^), and the mean light intensity controlled by the fraction of “on” time. With the period of 400 minutes, it results in minimal changes in mean gene expressions (Figure 1A), but a significant reduction of gene expression noise across the intermediate mean light intensities and intermediate mean gene expression level (purple lines in Figure1B-C). For the intermediate mean light intensities, the distribution spreading is reduced by “eliminating” the “low” and “high” states and generated an intermediate peak, which increases with mean light intensity (dashed line in Figure 1D). When the period of PWM shorten, the noise levels (Figure 1, B-C) and histograms gradually returned towards the AM scenario (Figure 1-figure supplement 2B-F). Therefore, we could independently modulate gene expression level and noise by altering the mean light intensity and period of PWM within the range sandwiched between blue and purple lines in Figure 1, B-C. A preliminary lookup table is computed by fitting expression level and distribution spreading (quantified as the ratio of mRuby at 90 and 10 percentiles) (Figure 1-figure supplement 2H, I). Such manipulation could be utilized in studying cell fate control.

To confirm the discovery, we stably transfected HeLa and F9 mouse embryonal carcinoma cell with a lightON-GFP circuit (plasmids A2, B2 in Figure 1-figure supplement 1B) to obtain HeLa-AB2 and F9-AB2 clones. We observed similar phenomena of large noise with AM and reduced noise with PWM with HeLa-AB2 (Figure 1-figure supplement 3A-C) and F9-AB2 (Figures-figure supplement 3D-K) cell. It suggested that the PWM-modulation of lightON gene expression noise could be ubiquitous in mammalian cell.

### Histone acetylation and transcription-epigenetic bistability

Previous studies of histone modification kinetics in CHO cell suggested that histone deacetylase 4 (HDAC4) could be related to our observed gene noise reduction, as its rate constants are the closest to the 400 minutes of PWM (Bintu et al., 2016). To identify histone modifications that contributed to the gene noises, we tested the effects of LMK-235, a selective inhibitor for HDAC4/5 and H3K9/H3K27 Deacetylation (Marek et al., 2013). Besides, we tested Vorinostat, a broad-spectrum inhibitor for HDACs (Finnin et al., 1999), and PF-06726304, a selective inhibitor for the Polycomb Repressive Complex 2 component and H3K27 trimethylation (Kung et al., 2016). We added them to the HeLa-AB1 cell and performed light inductions for 2 days with AM at 25 μW/cm^2^ before flow cytometry analysis. Only LMK-235 significantly increases the expression of mRuby, shown in Figure 2A-B. To compare mRuby expressions at the same GAVPO levels, we calculated the population average mRuby-*vs*-GFP-GAVPO response curves for cells treated with these inhibitors (Figure 2C). The mRuby-*vs*-GFP-GAVPO curve for LMK-235 treated cells (red) is significantly steeper than that of the control cells (blue), demonstrated that it increases explicitly mRuby expression for the cells with the same GAVPO level. Comparing the effects on AM and PWM with 400 min period, LMK-235 increases mRuby expression but has a minor impact on noise reduction (Figure 2-figure supplement 1A-F). LMK-235 also increase gene expression level for F9-AB2 cell and HeLa cell with Tet-On inducible expression system (Figure 2-figure supplement 2A-B).

The p65AD of the transcriptional activator GAVPO is reported to recruit CBP/p300, dual functional proteins that serve as both co-activators to recruit components of the mediator and transcriptional preinitiation complex (PIC), and as histone acetyltransferase (HAT) (Black et al., 2006; Gerritsen et al., 1997; Ogryzko et al., 1996). A recent CBP/p300 acetylome study reveals their rapid acetylation kinetics at many loci in mouse (Weinert et al., 2018). Put together, we propose a mechanistic model for transcription-histone acetylation coupling of the light-inducible expression systems (Figure 2D), (a) chromatin stochastically switching between silent and active states; (b) light induces GAVPO dimerization and enable its binding to UAS elements of the 5xUAS promoter in the active chromatin; (c) bound GAVPO recruits CBP/p300; (d) CBP/p300 acetylates nearby histones, and recruit PIC and mediator complex to facilitate transcription initiation; (e) the locus with high histone acetylation would tend to stay in the active state and further increase the chance of binding of GAVPO dimer. In summary, the DNA-GAVPO-HAT complex increases the probability of chromatin accessibility and GAVPO binding to DNA by increasing histone acetylation.

For AM illumination, this positive feedback loop could generate bistability, likely with intermediate GAVPO dimer concentration or light intensity (between cyan dashed line), illustrated in Figure 2E. HDACs deacetylate histone and contribute to set the boundaries between bistability and monostability. For some cells, the local promoter-GAVPO dimer interaction is high enough to initiate the positive feedback loop and lead to elevated local histone acetylation and high gene expression (red line). Other cells fail to start the positive feedback loop, which leads to low histone acetylation and transcription (blue line). Under this condition, the isogenic cell in a homogenous environment would exhibit large noise due to the history-dependent occupation of each state and stochastic switches between the two (Isaacs et al., 2003).

The principle for PWM-induced noise reduction could be that, the cell is switching between high monostable state (red) at 100 μW/cm^2^ and low monostable state (blue) at dark. In each state, the noise is low, and cell pass through the bistability region rather quickly. For PWM with shorter period, cell doesn’t have enough time to settle at either of two low-noise states, which leads to high noise. To test this, we used F9-AB2 cell (with higher dynamic range) and PWM with a period of 400 min with decreasing maximum light intensities. With maximum light intensity of 75 μW/cm^2^ or higher, PWM generates narrow histograms (Figure2 – figure supplement 2H-I). Yet with maximum light intensity of 50 μW/cm^2^ or lower, PWM lost the ability to reduce histogram dispersion (Figure2 – figure supplement 2J-L). This observation suggests that the boundary between bistability and high monostability is between 50 and 75 μW/cm^2^.

To further validate the mechanism, we constructed an ordinary differentiation equations (ODE) model (Figure 2D), with the details described in method section. The simulated histograms of mRuby with AM and PWM light inductions are shown in Figures 2F-G. It is consistent with the experimental observation that AM induced broad gene expression spreading at intermediate light intensities, which PWM with 400 min generated narrower spreading in mRuby expression. The *in silico* inhibition of HDAC activities shows the bistability remains with decreasing in the light threshold (Figure 2H), consistent with LMK-235 experiments (Figure 2A-B, Figure2 – supplement 2A). Furthermore, the model predicts that selective inhibition of HAT activities of CBP/p300 disrupt the existence of bistability (Figure 2I).

### Disruption of epigenetic bistability by static inhibiting HATs and dynamic alternating chromatin accessibility

We proposed that large gene expression noise is the consequence of epigenetic bistability induced by the positive feedback loop formed among promoter-bound GAVPO, HATs, and histone acetylation. We validated this prediction utilizing A-485, a highly selective and potent inhibitor for HAT activities of CBP/p300 (Lasko et al., 2017; Weinert et al., 2018) to disrupt the positive feedback loop without changing their transcriptional initiation function. When it was added to HeLa-AB1 cell under AM light inductions, the CV values for mRuby expressions exhibit significant reduction, especially at intermediate light intensities and mean mRuby expression levels (Figure 3B-C). It seems that A-485 eliminates the “low” and “high” expression states and generates an intermediate state with narrower mRuby expression dispersion (Figure 4D-E). It also confirms that CBP/p300 are the prominent co-activators involved in transcription and histone modifications. The mean mRuby expressions were also reduced (Figure 3A), as HDACs could tilt the balance towards lower histone acetylation, which leads to lower chromatin accessibility and less transcription. To validate this explanation, we added both A-485 and LMK-235 to the cell. As shown in Figure 3, A-C and F, additional HDAC4/5 inhibitor increases the expression level back to light induction control while retains the noise reduction phenomenon. In principle, simultaneous inhibition of CBP/p300 HAT activity and HDAC4/5 could also be used to reduces noise without changing mean gene expression. However, their effects on either synthetic and endogenous gene expression have never been examined.

**Figure 3.**
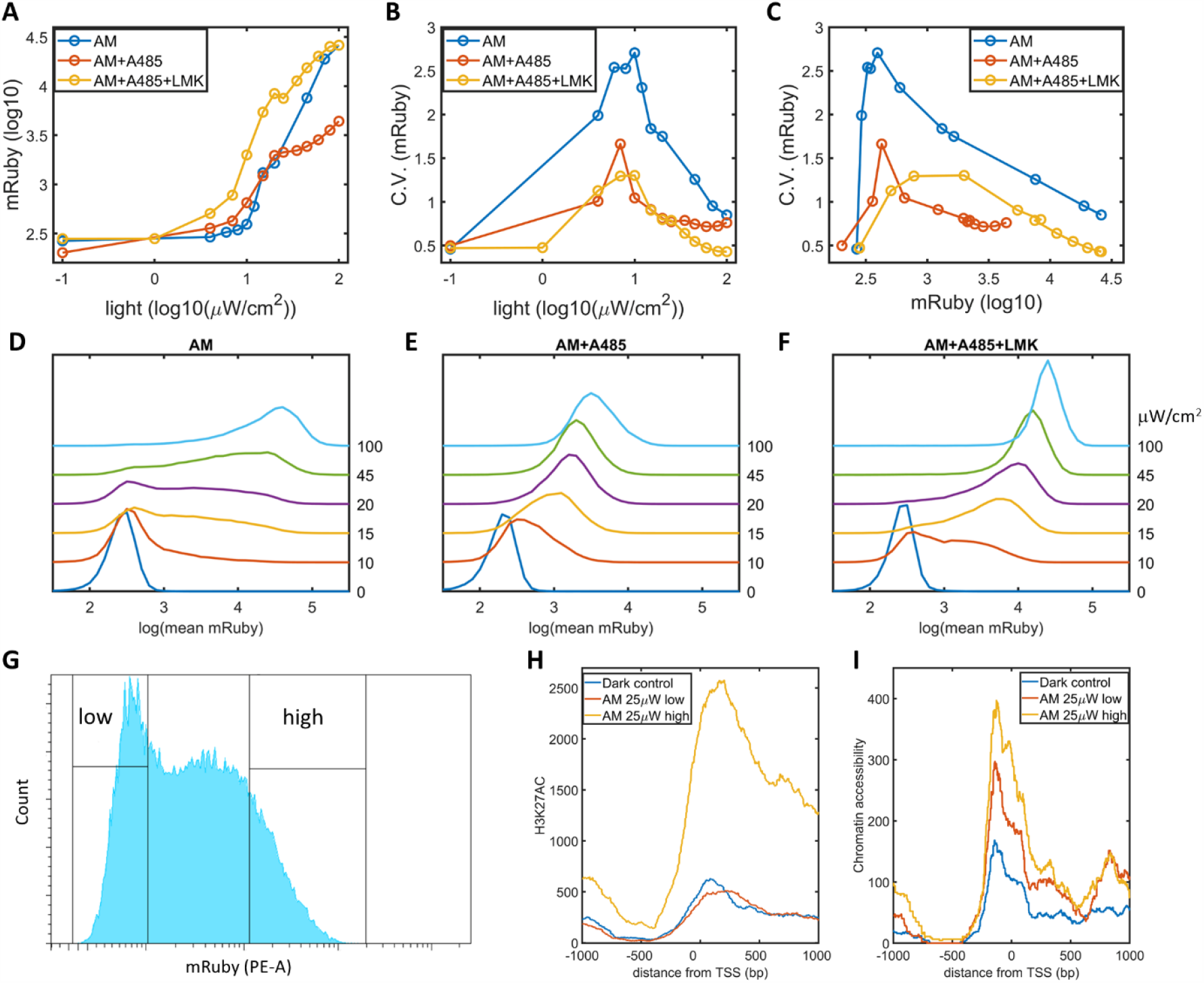
Validation of epigenetic bistability of HeLa-AB1 cells with CBP/p300 inhibitor, H3K27ac, and ATAC-seq analyses. (**A-C**) Dose-response curve of mean mRuby versus light intensities (A), CV of mRuby versus light intensities (B) or vs mean mRuby (C). The experimental conditions include AM modulation alone (blue), AM with 1 μM A-485 (red), and AM with 1 μM A-485 and 0.333 μM LMK-235 (yellow). (**D-F**) Histograms of mRuby at increasing light intensities for AM alone(D), AM with 1 μM A-485 (E), and AM with 1 μM A-485 and 0.333 μM LMK-235 (F). Specific light intensities are marked to the right of each histogram. (**G**) HeLa-AB1 cells treated for 1 day of AM light induction with 25 μW/cm^2^ were sorted into low- and high-mRuby expression populations for further ChIP-seq and ATAC-seq analysds. (**H-I**) H3K27ac ChiP-seq (H) and ATAC-seq (I) analyses for inserted 5xUAS promoter, aligned at TSS for dark control (blue), low-mRuby expression population from AM 25μW/cm^2^ (red), and high-mRuby expression population from AM 25μW/cm^2^ (yellow). Dark control is collected from cells kept in the dark. Each sample contains 10,000-50,000 cells.

**Figure 4.**
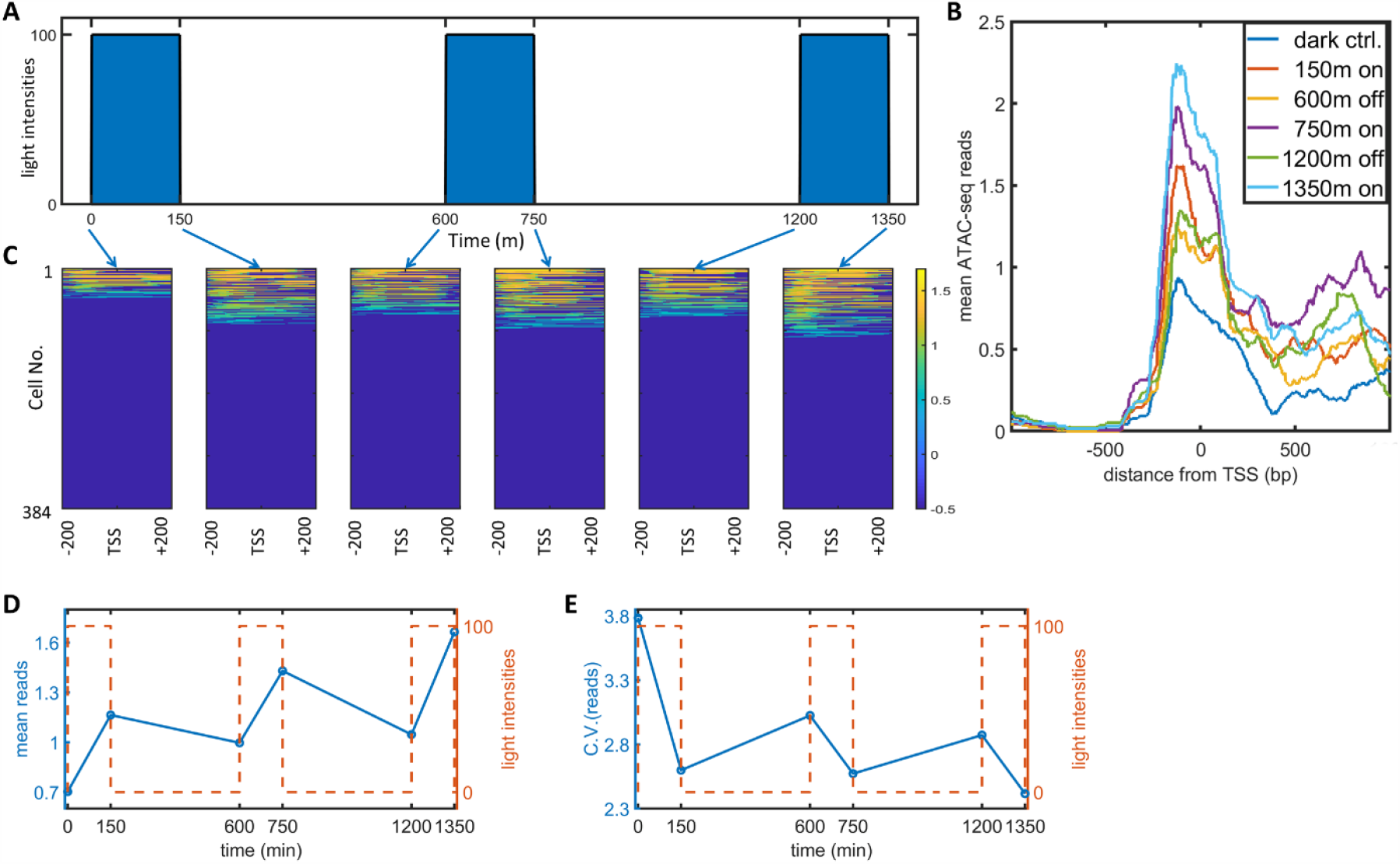
Single-cell ATAC-seq analysis of dynamic noise reductions for HeLa-AB1 cells with PWM. (**A**) Dynamic PWM induction program used to generate cells for single-cell ATAC-seq, and times for collections. As indicated with blue arrows, there are 6 populations of cells collected, at dark control (0 m), 1 light on (150 m), 1 light on-off cycle (600 m), 1 light on-off cycle and 1 light on (750 m), 2 light on-off cycles (1200 m), 2 light on-off cycles and 1 light on (1350 m). (**B**) The average ATAC-seq reds over single cells against distance from TSS of 5xUAS promoter for 6 cell populations. (**C**) Heatmap plots of single-cell ATAC-seq reads from −200 to 200bp around TSS of 5xUAS promoter. Reads are display in common logarithms. The x-axis represents the distance from TSS. The y-axis represents the cell index. Color represents the common logarithm of the reads. There are 384 cells for each time point. For each cell, a read was further defined as the average over 400 bp. (**D-E**) The mean reads (D) and CV of reads (E) over 384 cells were plotted against time. Red dashed lines represent the PWM light induction dynamics.

To directly assess the existence of epigenetic bistability of HeLa-AB1 cell, we used AM with 25 μW/cm^2^ light for 1 day to induce large expression dispersion, and sorted the mRuby-low and mRuby-high populations (Figure 3G). We performed ChIP-seq assays for H3K27ac and mapped the reads to the whole genome sequence assembly of this HeLa-AB1 clone containing lightON expression cassettes at 9 loci (Figure 3 – figure supplement 1A-B). The reads from these loci were combined using the transcriptional starting site (TSS) as the reference (Figure 3H). There is a peak spanning from the 5xUAS promoter (−200 bp) to the N-terminus of the mRuby gene (Figure 3 – figure supplement 1A). The signals for the mRuby-low and the dark control populations are essentially the same, and the signal for mRuby-high population is about 4 times higher. These demonstrate that the mRuby-high population from intermediate light induction is at a much higher epigenetic active state than the mRuby-low and dark control ones. We also performed ATAC-seq assay to the same cell populations. As shown in Figure 3I, the chromatin accessibility is higher at the closer vicinity of TSS, approximately from −200 to +200 bp. The hierarchy of chromatin accessibility is mRuby-high, mRuby-low, and dark control, with less separation between the high and low populations than H3K27ac, and clear difference between mRuby-low and dark control. Hence H3K27ac is related to lightON gene expression, consistent with the hypothesis of epigenetic positive feedback loop.

To observe the effect of PWM induction on chromatin accessibility variations, we performed single-cell ATAC-seq analysis of the dynamic process of PWM induced noise reduction. Specifically, we set up a PWM regiment with 100 μW/cm^2^ maximum light, 600 min period, and 25% on-ratio (150 min). We collect cells at 6 time-points corresponding to different light cycle stages, as illustrated in Figure 4A, and sorted 384 tagged nuclei for each population for sequencing and analysis. The mean reads spanning −200 to +200 bp exhibit graduate increases over each light induction cycle, shown in Figure 4B. The single-cell reads for the six populations are plotted as heatmap images (Figure 4C). Since the chromatin open-and-close is a dynamic process, it is feasible we observed at most approximately 30% cells with reads. Nevertheless, the fraction of cells with opening chromatin is lowest with dark control. Each cycle of “on” light (150, 750, 1350 min) leads to more cells with reads. At the end of each “on-off” cycle (600, 1200min), the fractions of cells with reads didn’t go all the way down to the level of dark control but increase over the cycles. To obtain quantitative measurements, we took the mean reads over 400bp for each cell and calculated the means and CVs for all populations, plotted against time (Figure 4D-E). The mean reads started at around 0.7 increases after each “on” period and decreased to a less extent after each “off’ period. In the meantime, the CV for the mean reads started with a high CV value (high noise of chromatin openness) of 3.8 and reduced drastically to 2.6 following just 150 minutes of illumination. It increases after each further “off” period and decreases after each further “on” period to less extents. These data indicated that the basal (dark) chromatin openness is very heterogeneous. A short period (150 min or less) of 100 μW/cm^2^ light not only initiates transcription but also reduces heterogeneity of chromatin openness. Each consequent “on” period further increases the chromatin openness and reduces the heterogeneity, and the “off’ period maintains part of the epigenetic “memory.” Therefore, the chromatin accessibility heterogeneity is reduced after each cycle of “on” and “off.” But the heterogeneity still exhibits temporal fluctuations as there is a difference between the ends of “on” and “off” states.

### RNA dynamics flattens PWM-induced pulsatile chromatin opening

To connect the observations of PWM-induced noise reduction between epigenetic level and protein level, we measured nuclear mRNA dynamics for single cells at AM and PWM light induction scenarios utilizing an MS2-MCP system (Tutucci et al., 2018). HeLa-AB2 cell, which exhibits PWM induced noise reduction at periods of 400 and 200 min (Figure 1 – figure supplement 3B-C), was further stably transfected the ligthON mRNA imaging circuit (Figure 1-figure supplement 1A, plasmid C2) to obtain HeLa-ABC2 cell. We used live-cell spinning disk confocal microscopy to image the single mRNA. When multiple MCP-tdTomato molecules bound to an mRNA containing 24 MS2 hairpins, they form a punctum with a diameter around 500 nm above the background fluorescent, as illustrated as single mRNA puncta in the two nuclei in Figure 5A. The single-cell mRNA counts for all the tracked cells are plotted against time as heatmap images for AM (100 μW/cm^2^), PWM (100 μW/cm^2^, 200 min, 25% on-ratio), and AM (25 μW/cm^2^), as shown in Figure 5B-D, respectively. For quantitative analysis, the CVs for single nuclear mRNA numbers are plotted against time (Figure 5E). For the AM with 100 μW/cm^2^, there variation is large at the beginning and reduced overtime to lower level (Figure 5, B, E). This is consistent with the time course flow cytometry analysis of mRuby protein (Figure 5F). For the AM with 25 μW/cm^2^, it exhibits higher variations over time (Figure 5, D, E). For the PWM case, there is a clear pulsatile nuclear mRNA counts with a period of 200 minutes, but mRNA appeared with a time delay of about 40 minutes after each “on” time point, and they remain in the nuclear for the next 100 minutes or so before translocated to the cytoplasm (Figure 5C). Two phenomena contribute to noise reduction by PWM. First of all, the appearance of nuclear mRNA (transcription) exhibits stochasticity consistently much lower than the AM with 25 μW/cm^2^, no matter if the cells were in the light “on” or “off” periods (Figure 5E). This suggested that the cells were likely alternating between high and low epigenetic states and not trapped in the bistability region (Figure 2D). Secondly, 50 minutes of light pulses lead to nuclear mRNA pulses with wider durations (∼100 minutes) and delays (∼40 minutes). The mRNA transcription delay, motion, and transportation to cytoplasm effectively “flatten” the pulsatile curve and filtered out the temporal “noise” at chromatin accessibility and transcriptional level. It would be further reduced at the protein level with several the downstream accumulation mechanisms such as the longer half-life of protein *etc*.

**Figure 5.**
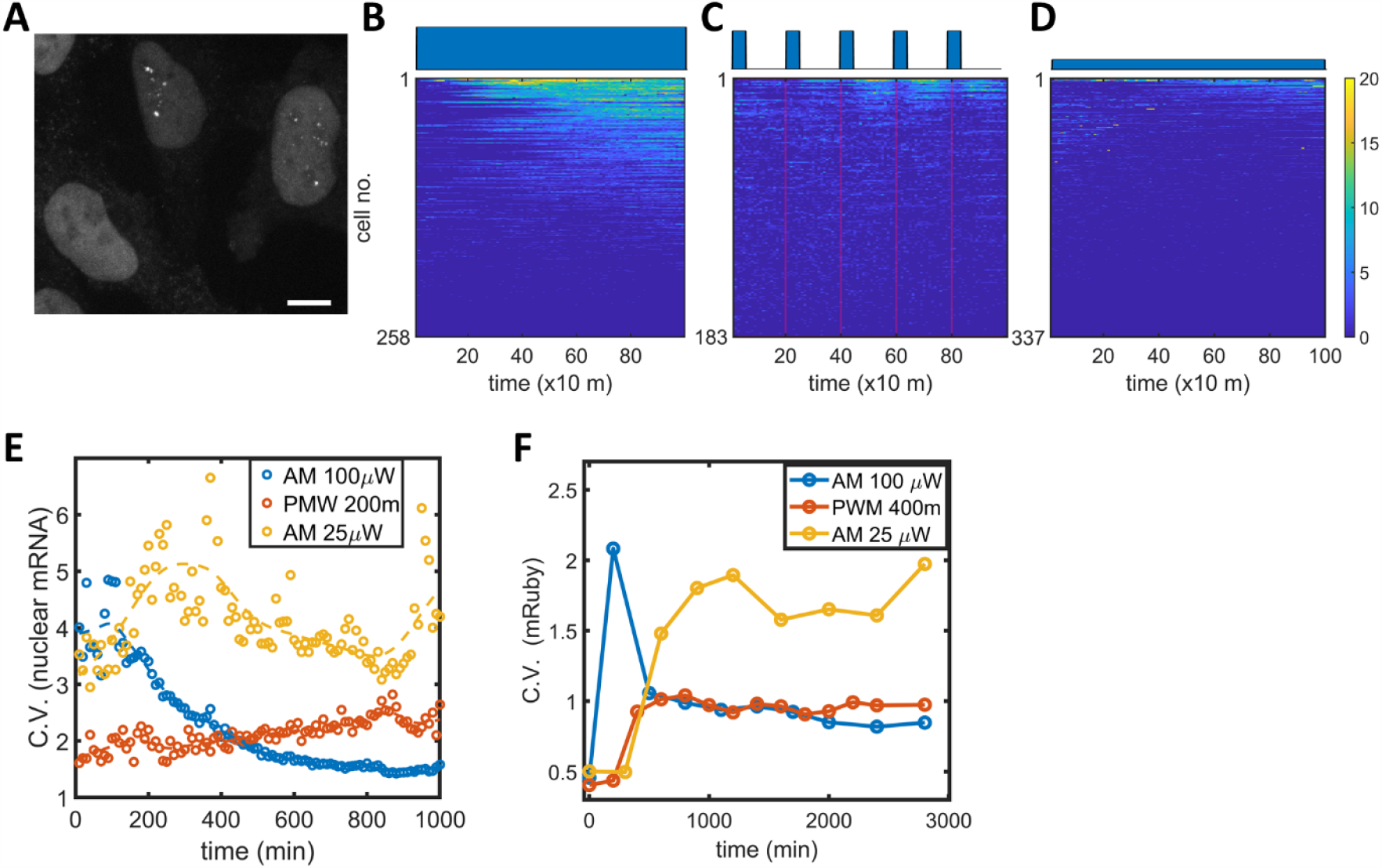
Single mRNA live-cell imaging for HeLa-ABC2 cells with AM and PWM. **(A**) A representative z-projection image at one time point. The bright blob shapes (diameter around 10 μm) represent the nuclei. The small brighter puncta (approximately 500 nm in diameter) in the nuclei represent nuclear mRNA molecules. The darker fluorescent signals are cytosol fluorescent signals neglected in this analysis. The scale bar represents 10 μm. (**B-D**) The single-cell mRNA counts over time are detected and shown as heatmaps for AM with 100 μW/cm^2^(B), PWM with a maximum of 100 μW/cm^2^, on-fraction of 0.25, and a period of 200 m (C), and AM with 25 μW/cm^2^(D). The x-axis represents time with a unit of 10 minutes. The y-axis represents the cell index. The filled dark blue plots on top of heatmap images represent the light induction schemes. (**E**) The CV for single-cell nuclear mRNA plotted against time for AM with 100 μW/cm^2^ (blue), PWM with a period of 200 m and an on-fraction of 0.25 (red), and AM with 25 μW/cm^2^ (yellow). (**F**) The CV of mRuby protein at different time points for AM with 100 μW/cm^2^ (blue), PWM with a maximum intensity of 100 μW/cm^2^, a period of 400 m and an on-fraction of 0.25 (red), and AM with 25 μW/cm^2^ (yellow).

### Evidences of dual-functional CBP/p300-induced heterogeneity in endogenous genes

CBP/p300 is prominently implicated in regulation of enhancer-dependent cell-type-specific gene regulations (Ogryzko et al., 1996; Weinert et al., 2018). Compiled analysis of vast public ChIP-seq data suggests more than ten thousand potential CBP/p300 targeted genes from both human and mouse cell experiments (Oki et al., 2018) (https://chip-atlas.org). Most of the targeted genes have complexed gene regulations to overshadow the effects of potential positive feedback loop formed by transcriptional and histone acetylation functions of CBP/p300. Nevertheless, this positive feedback loop by CBP/p300 could potentially increases heterogeneity in a subset of endogenous genes. To find evidence that positive feedback loop by CBP/p300 could upregulated heterogeneity in endogenous genes, we use the A-485 and LMK-235 to inhibit CBP/p300 and HDAC4/5 to disrupt the potential positive feedback. In mouse ES cell, p300 is recruited by the master transcriptional factors Nanog, Oct4, and Sox2 to facilitate ESC-specific gene expressions (Chen et al., 2008). Mouse ES cell cultured with LIF (but not 2i cocktail) exhibits subpopulations with high and low expression level of Rex1, Nanog, and dynamic transitions between the two “states” (Singer et al., 2014; Toyooka et al., 2008). To test whether the CBP/p300 positive feedback loop could contribute to the “bistability” in mouse ES cell, we constructed a Rex1-GFP/Oct4-mCherry Knock-in cell line with D3 cell. It exhibits bimodal Rex1-GFP expression, with 97% high and 3% low subpopulations, and unimodal Oct4-mCherry expression (Figure 6A). After treatment with LMK-235 and A-485 for two days, the Rex1-GFP expressions converged to a unimodal expression centered between the high and low states (Figure 6B), like the HeLa-AB1 cell which treated with similar concentration inhibitors and AM light inductions (Figure 3D and F). Although inhibition of CBP/p300 globally also invokes indirect actions to Rex1 expression, this result suggests that it is possible that CBP/p300 could contribute to mouse ES cell heterogeneity by forming positive feedback loop of transcription initiation and histone acetylation.

**Figure 6.**
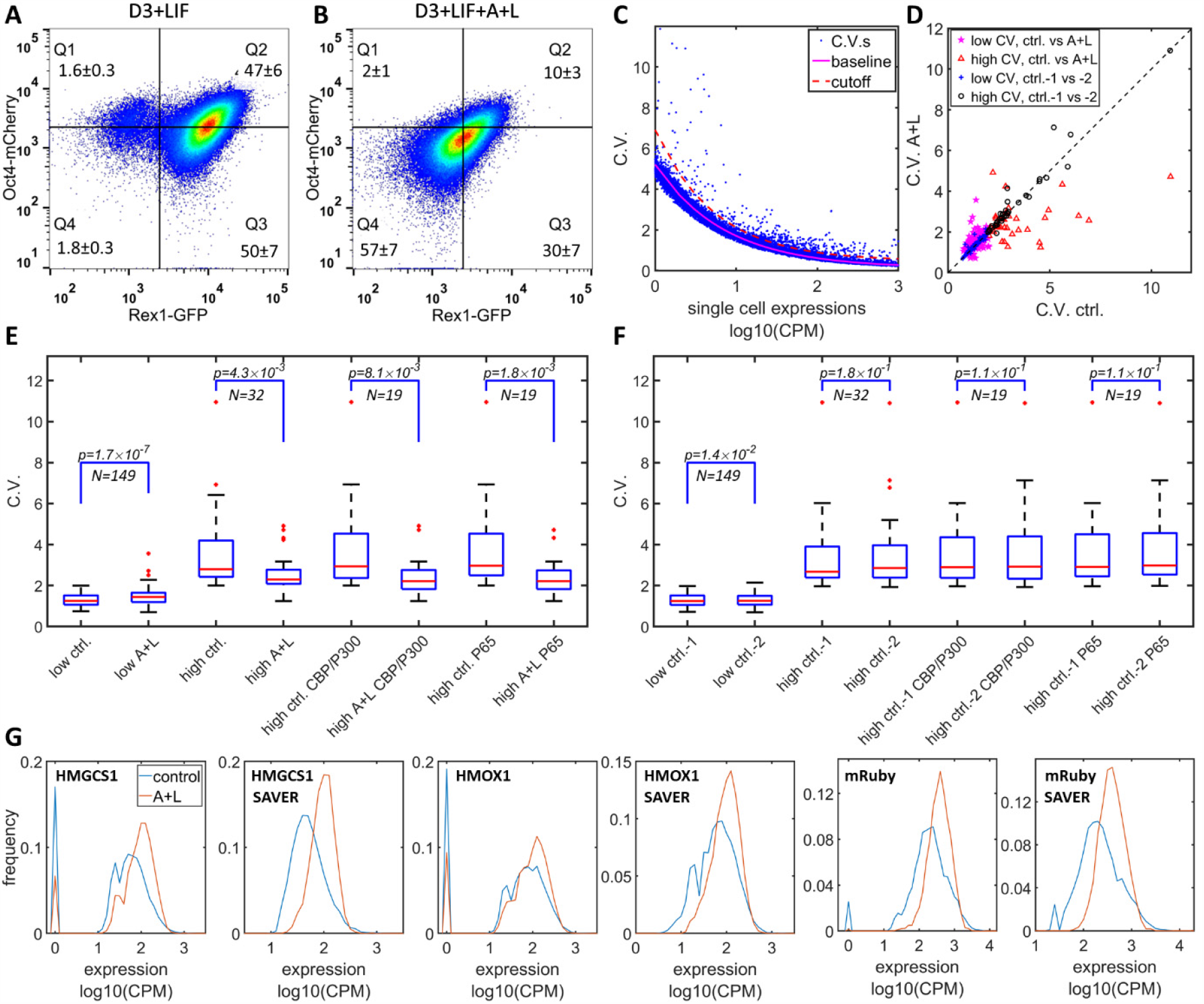
Simultaneously inhibition of CBP/p300 and HDAC4/5 reduces heterogeneity in endogenous gene expressions in mES D3 and HeLa cells. (**A, B**) The density plots of Oct4-mCherry vs Rex1-GFP for Mouse ES D3 double knock-in cell line with regular mESC medium (A), or with addition of 0.5 μM LMK-235 and 1 μM A-485 for 2 days (B). The mean+s.d. for the percentages of cells in Q1-Q4 were calculated from 3 (A) and 6 (B) biological repeats. (**C**) The CV vs mean expressions plot with scRNA-seq data from HeLa cell with or without 0.333 μM LMK and 1 μM A-485. Purple solid line represents the fitted power function of all the genes, with exponent of −0.48. Red dashed line represents the 3-sigma cut off for noisy outlier genes. (**D**) The genes with CV vs gene expression above the cutoff (red line in C), and mean reads >5 CPM were selected, and plotted as CV between HeLa cell with A485+LMK-235 and control condition. Purple pentastars and red triangles represent the genes with low (< 2.0) and high CV (≥2.0), respectively. Blue asterisks and black circles represent the CV of the same gene sets between two biological replicates of control cells. (**E**) Statistical analysis of the CV of genes between A485+LMK-235 and control, with low CV, high CV, high CV and positive for CBP/p300 or p65 ChIP. The boxes show the lower and upper quartiles, the whiskers show the minimum and maximal values excluding outliers; the line inside the box indicates the median; outliers (red dots) were calculated as values greater or lower than 1.5 times the interquartile range. The p values calculated using paired student *t*-test is shown in the figure. (**F**) Similar statistical analysis of CV between two biological replicates of control conditions for the same sets of genes. (**G**) Histograms of expressions for control (blue) and A485+LMK-235 group (red) of genes HMGCS1, HMOX1 and exogenous mRuby, calculated from normalized data without or with SAVER denoising algorithm. The peaks at “0” in the histograms represent cells with zero reads.

We further examined the effects of CBP/p300 and HDAC4/5 inhibitions on global gene expression heterogeneity in HeLa-AB1 and F9-AB2 cells by performing scRNA-seq analysis. HeLa-AB1 cell was either treated with LMK-235 and A-485 and blue light for two days (A+L) or light only (control). The detailed library construction, sequencing and data process are described in the method section. As shown in Figure 6C, when plot the CV vs mean expression, most of the genes concentrated on a narrow band, fit to a power function with exponent of −0.48 (solid magenta line), similar with the predicted exponent of −0.5 as a Poisson distribution. Outlier genes significantly deviate from this band (blue dots above the dashed red line) are further selected for assessment of the effects of A-485 and LMK-235. For the filtered gene set with lower CV values (less than 2 in control samples), there is a slight but statistically significant increase in mean CV from 1.30 to 1.45, as shown in Figure 6D (magenta pentastars) and E. On the other hand, for the filtered gene set with higher CV, there is clearly much more genes with large reductions in CV with the inhibitions of both CBP/p300 and HDAC4/5, shown in Figure 6D (red triangles). For this set of genes, the mean CV exhibits a large (3.49 to 2.55) and significant reduction. Within this set, targeted genes by CBP/p300 or p65 show similar mean CV reduction, shown in Figure 6E. There is much less deviation between the two biological repeats of control samples, especially for the high CV gene sets, as shown in Figure 6D and F. The reduction in CV can be visualized with narrower histograms by inhibition of both CBP/p300 and HDAC4/5, computed from UMI counts, without or with SAVER denoising process (Huang et al., 2018; Luecken and Theis, 2019), shown in Figure 6G. Similar phenomena was observed with F9-AB2 cell, shown in Figure 6-figure supplement 1. These results provide first indirect evidences that CBP/p300 positively contribute to global gene expression noises, especially for highly heterogenous genes in human and mouse cells. It could also contribute to the heterogeneous states in mouse embryonic stem cell.

## Discussion

Synthetic inducible gene expression systems were developed to study the function of genes in various cellular and physiological processes. The widely used ones enable us with the practicality of induction signals such as doxycycline and light, and high dynamic range of expression, *etc*. They are essentially the simplest form of gene regulation involving one transcriptional activator. However, they often exhibit huge gene expression noises, especially in mammalian cell (CV as high as 5-10, shown in Figure 1 - figure supplement 3 and Figure 2-figure supplement 2C). The main molecular mechanism that contributes to such large noise has not been identified. In this study, we performed a quantitative characterization of gene expression noise in a light-induced expression circuit (Wang et al., 2012) in human and mouse cell under amplitude modulation and pulse width modulation of light induction. We found that PWM light controls noise in a period-dependent manner, which enables a phenomenological approach for independent modulation of expression level and gene expression dispersion with proper mean light intensity and period of the PWM. It would support more precise designs of cell fate control studies. Hardwired negative feedback circuits could reduce gene expression noise in mammalian cells, but cannot independently modulate noise and expression (Guinn and Balázsi, 2019; Nevozhay et al., 2013).

Based on the overlapping time scales of PWM noise reduction and epigenetic regulations by HDAC4 and CBP/p300 (Bintu et al., 2016; Weinert et al., 2018), we identified a positive feedback loop in transcription and histone regulation. It is formed with light-induced GAVPO dimer binding to 5xUAS promoter in transiently opened chromatin, its p65 activation domain recruits CBP/p300 to facilitate transcription initiation and histone acetylation, and acetylated histones to increase chromatin accessibility and GAVPO binding. This positive feedback loop is predicted to generate bistability and large gene expression noises at intermediated light inductions at intermediated light inductions. PWM reduces noise by alternating the cell between two monostable states. This mechanism is experimentally validated by disruption of bistability with selective inhibitor A485 for CBP/p300 HAT activity, identification of low and high states of H3K27AC, and chromatin accessibility in the low and high expression cell populations in the bistability region.

We further use single-cell ATAC-seq analysis and live-cell single mRNA imaging to illustrate the dynamic process of noise reduction for PWM noise reductions. The scATAC-seq unveils that the chromatin accessibility is highly variable at dark control and reduced after each cycle of light induction. Nevertheless, there is a temporal fluctuation of the chromatin accessibility within each period. The noise level of single-cell nuclear mRNA counts for PWM remains lower than the AM with intermediate light. Similar phenomena were observed at the protein level. To sum it up, the PWM noise reduction is effective at the epigenetic level by switching between the low and high monostable regions and avoiding the bistability region. The fluctuations from pulsatile chromatin accessibility and transcription events are averaged down at nuclear mRNA level through accumulation and further filtered at protein level due to longer protein half-life.

CBP/p300 are reported to targeted large sets of genes and primarily regulate enhancer-dependent cell-type-specific gene expressions. We tested the effect of potential disruption of this positive feedback loop on endogenous gene expression using both A485 and LMK-235. In a mouse ES-D3 Rex1-GFP reporter cell line treated with both inhibitors, we found the Rex1-high and Rex1-low subpopulations disappeared and resulted in a single population with intermediate Rex1 expression. In HeLa-AB1 cell and to a less extend in F9-AB2 cell, scRNA-seq analysis suggested that the two inhibitors preferentially reduced the heterogeneity of high noise genes. These preliminary results suggested that the transcription-epigenetic positive feedback loop involved in CBP/p300 could contribute to the highly heterogeneous endogenous gene in mammalian cells.

In summary, we identified a transcription-epigenetic positive feedback loop involving p65AD and CBP/p300, which could lead to bistability and large gene expression noise. Since p65AD and VP16, used in most synthetic gene expressions and many endogenous gene regulations (Gilbert et al., 2013; Gossen and Bujard, 1992; Gossen et al., 1995; Khalil et al., 2012; Wang et al., 2012), are all reported to interaction CBP/p300 and other HATs (Black et al., 2006; Gerritsen et al., 1997; Goodman and Smolik, 2000; Kim et al., 2012; Wang et al., 2000), this could be a general mechanism for modulation gene expression noises in mammals. CBP/p300 could also contribute to the noise in some of the endogenous genes in a similar manner. We found that pulse light induction with a long period could reduce noise as it helps the cell avoiding staying in the bistable regime. A picture of the dynamic process of this noise modulation has been established from chromatin accessibility to mRNA and protein levels using single-cell analysis. The noise reduction exhibits dose-response to the period of PWM, which enables independent modulation of mean gene expression and noise in the future quantitative study of cell fate control.

## METHODS DETAILS

### KEY RESOURCES TABLE

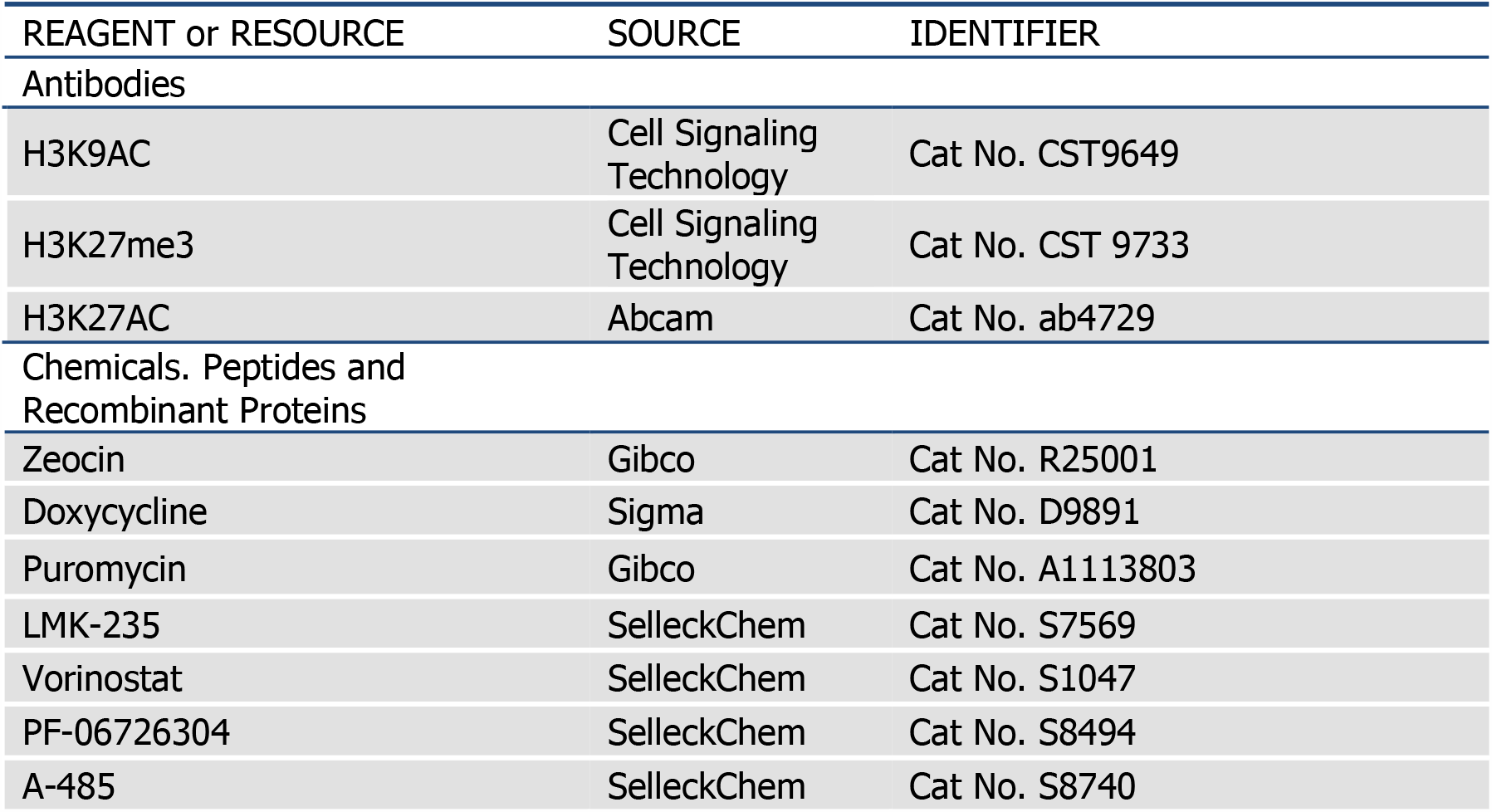
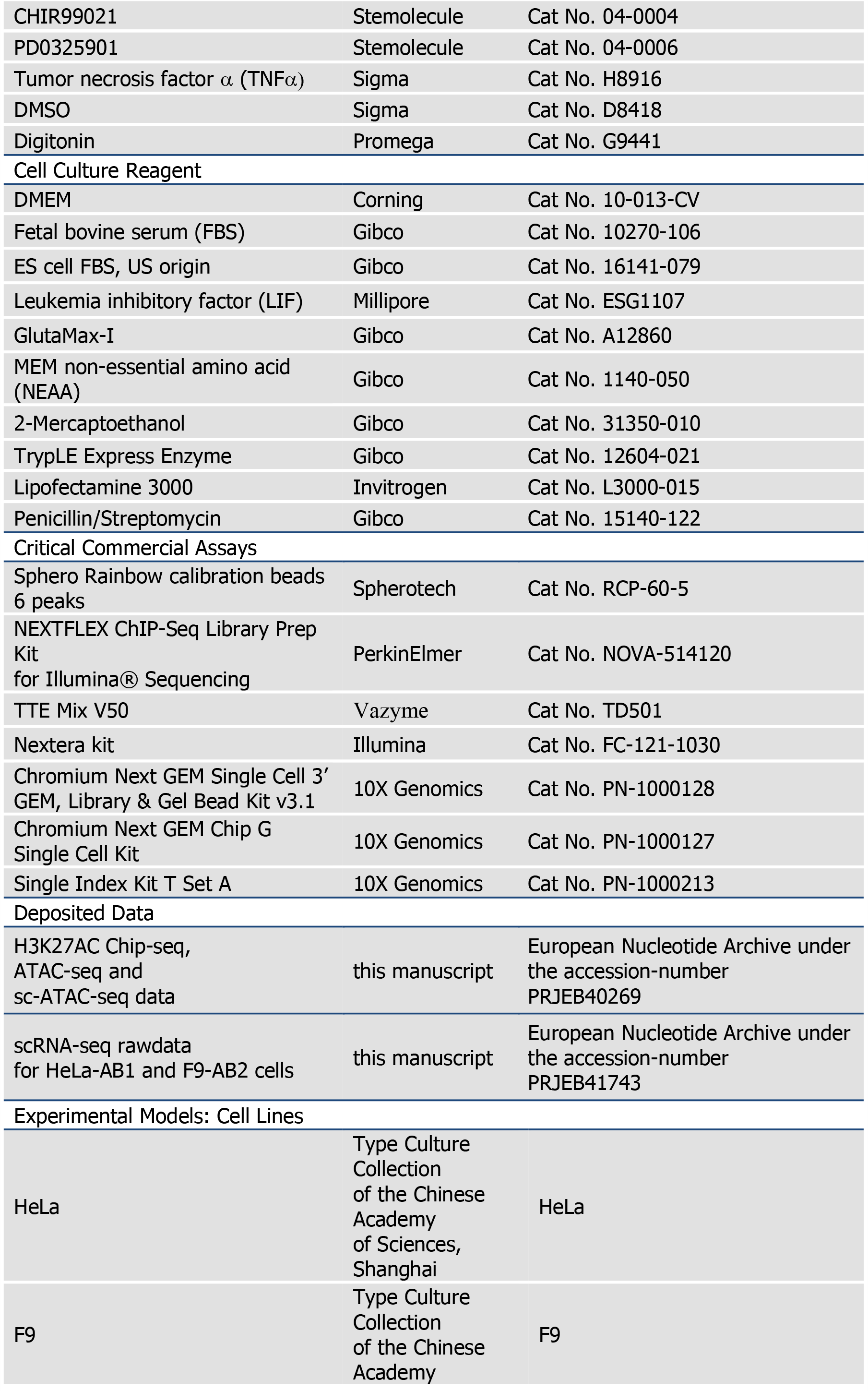
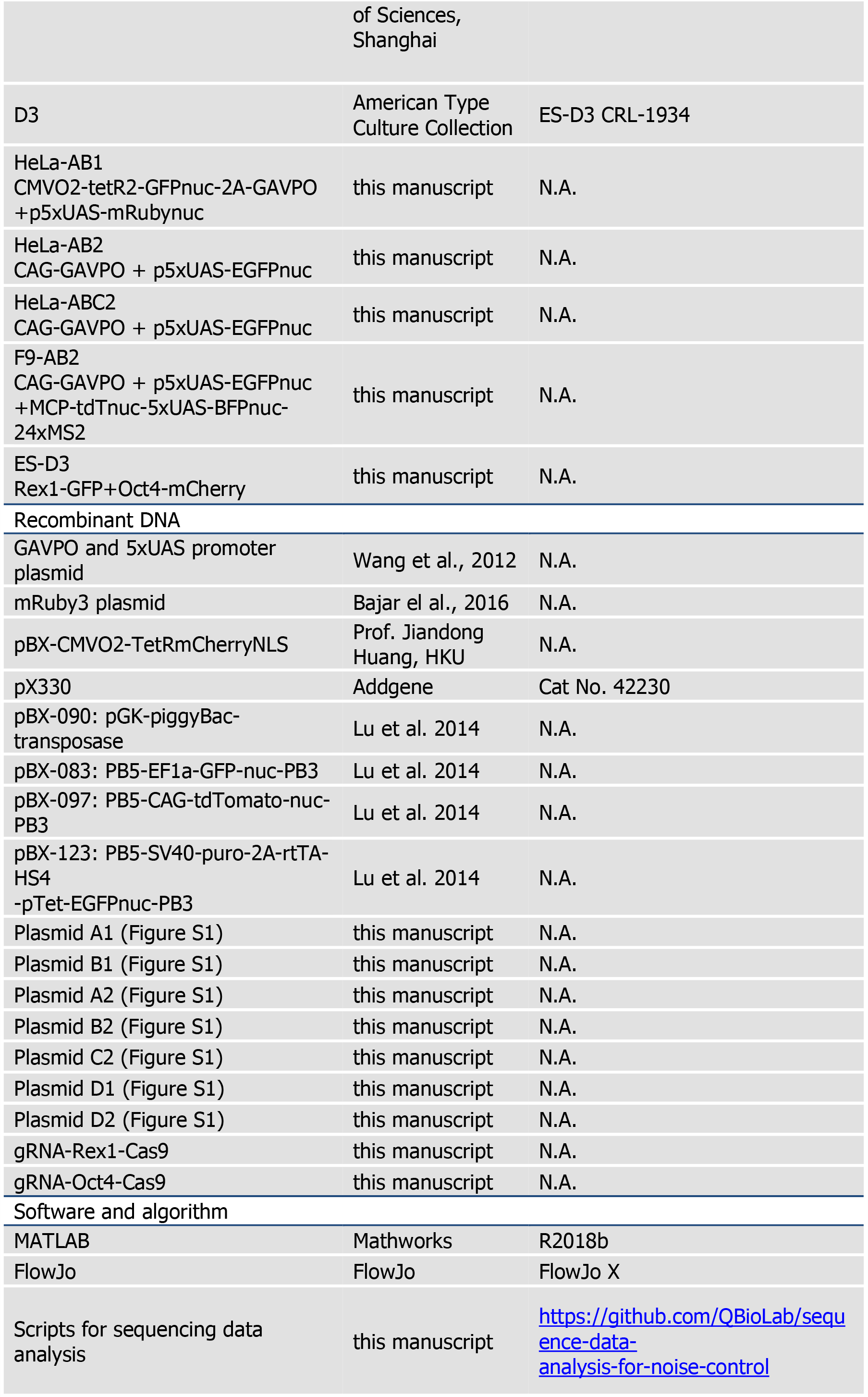
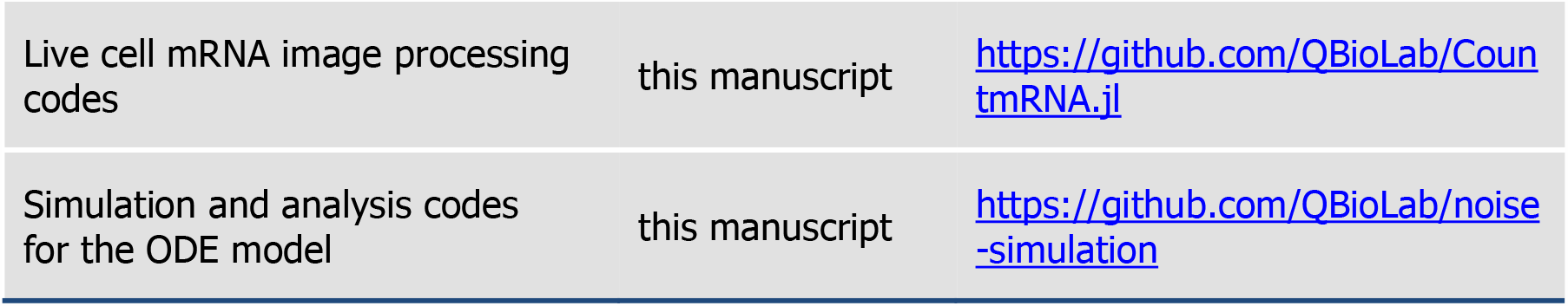

### Plasmid constructions

Every version of the lightON system is constructed with two plasmids, a plasmid A with a promoter-driven expression of the synthetic light-sensing transcriptional activator GAVPO, a fusion protein of GAL4 DNA binding domain, engineered light-sensing VVD domain and p65AD activation domain, and a plasmid B with ligthON promoter (5xUAS) driven reporter fluorescent proteins, (Figure S1B). The plasmids containing 5xUAS promoter and GAVPO (Wang et al., 2012) were gifts from prof. Yi Yang. The plasmid for tetR negative feedback circuit similar to (Nevozhay et al., 2013), containing pCMV-tetO2-tetR-GFP-nuc was a gift from prof. Jiandong Huang. The plasmid containing mRuby3 was a gift from prof. Jun Chu. For the HeLa cell line used in flow cytometry, live-cell imaging and epigenetic assays, the transcriptional activator expression plasmid A1 is constructed by subcloning pCMV-tetO2-tetR-GFP-nuc with GAVPO, and a plasmid with PiggyBac transposon backbone (Lu and Huang, 2014) to form PB5-pCMV-tetO2-tetR-GFP-nuc-2A-GAVPO-2A-Zeo^r^-bGpA-PB3, and the lightON plasmid B1 is constructed by assembling 5xUAS, mRuby3, and subcloning into a plasmid with PiggyBac transposon backbone (Lu and Huang, 2014)to form PB5-5xUAS-mRuby-nuc-2A-Bla^r^-βGpA-PB3. For F9 and HeLa, the transcriptional activator plasmid A2 is constructed by subcloning GAVPO into another PiggyBac backbones to form PB5-pCAG-GAVPO-2A-Zeo^r^-βGpA-PB3, and the lightON plasmid B2 is constructed by subcloning 5xUAS, GFP-nuc into PiggyBac backbone plasmids to form PB5-5xUAS-GFP-nuc-2A-Bla^r^-βGpA-PB3. For the HeLa cell line used in mRNA imaging, in addition to transcriptional activator plasmid A2, lightON plasmids B2, one additional plasmid C2 was constructed by subcloning 5xUAS-tagBFP-nuc, 24 MS2 hairpin repeats (synthesized by Wuxi Qinglan Biotech), SV40 promoter, MCP-tdTomato-nuc (synthesized by Wuxi Qinglan Biotech) into a PiggyBac backbone plasmid to form PB5-pSV40-MCP-tdTomato-nuc-2A-Puro^r^-SV40pA-HS4-5xUAS-BFP-nuc-24xMS2-βGpA-PB3. In all these constructs, 2A stands for a 2A “self-cleaving” peptide T2A, derived from thosea asigna virus 2A, enable the same levels of transcription and translation for the two proteins. The nuc tag is a three tandem repeats of SV40 nuclear localization sequence facilitating the localization of the protein into the nucleus. The UAS is the specific GAL4 binding elements. The Puro^r^, Zeo^r^, and Bla^r^ are resistant genes for antibiotics puromycin, zeocin, and blasticidin, respectively. The HS4 is the Chicken hypersensitive site 4 (cHS4) insulator. The components of MS2-MCP system, MS2 RNA loop, and MS2 coating protein were adapted from ref. (Tutucci et al., 2018).

To generate knock-in Rex1-GFP and Oct4-mCherry mouse ES cell line, we constructed two donor plasmids D1 and D2, shown in Supplementary Figure 1. For Rex1-GFP knock-in donor plasmids D1, A T2A-GFP-nuc cassette is in-frame fused with an 5’ 1000 bp homologous arm including the C-terminal of Rex1 gene, and an 1000 bp 3’ arm including 3’UTR sequence of the Rex1 (Zfp42) gene. For the Oct4-GFP knock-in donor plasmid D2, an ATG-mCherry-linker cassette is in frame fused with an 800 bp 3’ arm including the N-terminal of Oct4 coding sequence, and an 800 bp 5’ arm including the 5’UTR sequence of Oct4. The gRNA sequences for Rex1 and Oct4, TCCTAACCCACGCAAAGGCC and CAGGTGTCCAGCCATGGGGA, respectively, were cloned into pX330 gRNA-Cas9 plasmid (Zhang Feng lab) to obtain Oct4-gRNA-Cas9 and Rex1-gRNA-Cas9 plasmids. All other components, otherwise specified, are adapted from (Lu and Huang, 2014). All the plasmid constructions were confirmed with Sanger sequencing. The sequences of plasmids and oligos used for the constructions will be provided upon request.

### Cell line constructions

To generate stable transfected HeLa cell with GFP-GAVPO and lightON-mRuby, HeLa cell is cotransfected with 400 ng plasmid A1, 400 ng plasmid B1, and 200 ng PGK-transposase plasmids using Lipofectamine 3000 (Catalog no. L3000-015; Invitrogen) following manufacture protocol in one well in a 24-well plate, and selected with 100 μg/mL zeocin (Catalog no. R25001; Gibco) for 6 days one day after transfection. The medium was replaced with DMEM supplemented with 10% FBS and 1μg/mL doxycycline (Catalog no. D9891; Sigma), and the cells were illuminated with blue light (100 μW/cm^2^) for 48h. Single-cell expressing both EGFP and mRuby were sorted to 96-well plates using BD FACSAria SORP. Each clone was verified using flow cytometry two weeks later. One clone, named HeLa-AB1 was chosen for further analysis. Similarly, F9 and HeLa cells were stably transfected with 400ng plasmid A2, 400 ng plasmid B2, and 200 ng PGK-transposase plasmids. The cells were illuminated with blue light for 2 days before sorting single cells with GFP expression. The difference is that single clones of F9 cells were analyzed 10 days after sorting. One F9 clone, named F9-AB2 was chosen for further analysis. Another HeLa clone, namely HeLa-AB2 was further transfected with mRNA imaging plasmid C2, and pGK-transposase plasmids, and selected with 1.5μg/mL puromycin (Catalog no. A1113803; Gibco) for 3 days. Cells were illuminated for 2 days with blue light and sorted with positive GFP, tdTomato, and BFP expression. One clone, namely, HeLa-ABC2 is chosen for single mRNA imaging study. To generate stable transfected HeLa cells with pTet-GFP, HeLa cell was further transfected with pBX-123 plasmid C2, and pGK-transposase plasmids, and selected with 1.5μg/mL puromycin (Catalog no. A1113803; Gibco) for 3 days. One clone, named HeLa-Tet-On was chosen for further analysis.

### Knock-in mouse ES cell line constructions

To generate knock-in Rex1-GFP and Oct4-mCherry mouse ES cell line, mouse ES cell line D3 (a gift from prof. Jianbo Yue) was cultured on 0.1% gelatin-coated 24-well plate with mESC medium (DMEM containing 15% ES FBS, 1000 U/ml LIF, 1X GlutaMax, 1X non-essential amino acid (NEAA) and 0.1 mM 2-Mercaptoethanol) until it reaches 50% confluence. D3 cells were transfected transfect with 500 ng Oct4-GFP donor plasmid (D1) and 500 ng Oct4-gRNA-Cas9 plasmids with Lipofectamine 3000, and diluted 1 day later to 6-well plate, and maintained for 4 days with the mESC medium containing 2i cocktail (3 μM CHIR99021 and 1 μM PD0325901). Single cells expression mCherry were sorted, expanded and genotyped with PCR and sequencing. One validated Oct4-mCherry clone was used to repeat the knock-in experiments with the Rex1-GFP donor plasmid (D2) and Rex-gRNA-Cas9 plasmid.

### Light induction expression experiment

HeLa-AB1 cell were plated in ibidi 24-well μ-plate at a density of 3×10^4^ cells/well. The medium was supplemented with 1 μg/mL doxycycline on day −1. On day 0, the dox-containing medium was replenished and the μ-plate was mounted to the 24-channel illumination apparatus located in a Thermo Heracell 150i CO2 incubator. Water cooling pump is turned on to ensure cells cultured at 37°C. A csv file containing experimental designed illumination intensities and dynamics for the 24 wells was loaded into the custom Python code to control the apparatus for 48 hours unless otherwise specified. For F9-AB2 and HeLa-AB2 cells, the procedure is similar except that doxycycline was not added.

### Finding PWM parameters to specific mean expression levels and distribution spreading

We took the flow cytometry data for HeLa-AB1 cell, with AM (period = 0) and PWM with different periods (such as Figure 1-figure supplement 2A-F), calculated the mean mRuby to mean light intensity curve, and distribution spreading (defined as the ratio of mRuby at 90 and 10 percentile) to mean light intensity curve. We then fitted the data using cubic smoothing spline function (csaps) with MATLAB 2018b to generate response surface plots for common logarithms of mean value, distribution spreading against period, and mean light intensities, shown in Figure 1-figure supplement 2H=I. We chose distribution spreading ratios, as it provides a more intuitive picture of how diverse the gene expressions instead of the coefficient of variations. Specifically, it is a measurement of how much difference in expression levels in the top 10 percentile to lowest 10 percentile, invariant to instrumentation settings.

### Experiments with inhibitors of epigenetic regulators

HeLa-AB1 cell were plated in ibidi μ-plate 24-well plate at a density of 3×10^4^ cells/well. The medium was supplemented with 1 μg/mL doxycycline on day −1. On day 0, the medium supplemented with 0.333 μM (unless otherwise specified) LMK-235 (Catalog no. S7569; SelleckChem), 0.333 μM Vorinostat (Catalog no. S1047; SelleckChem), 0.333 μM PF-06726304 (Catalog no. S8494; SelleckChem), or 1 μM of A-485 (Catalog no. S8740; SelleckChem). On day 0, the μ-plate was mounted to the 24-channel illumination apparatus located in the CO2 incubator and illuminated for 2 days before flow cytometry analysis. For Oct4-mCherry-Rex1-GFP mES cell, 2i cocktail was removed from mESC medium at least 4 days earlier to establish heterogenous mES cell culture with Rex1-GFP-high and Rex1-GFP-low populations. Cells were then treated with 0.5 μM LMK-235 and 1 μM A-485 for 2 days before performing flow cytometry analysis.

### Flow cytometry analysis

Cells in a 24-well were monodispersed using 125 μL of 1X TrypLE™ Express Enzyme (Gibco), neutralized with 250 μL of culture medium, filtered through 40 μm cell strainer (BD Falcon) to remove clumps, and analyzed using a Beckman Cytoflex S cytometer (Brea, US) equipped with 405/488/561/640 nm lasers at high speed (60 μL/min) for 2 minutes per sample. The rest of the samples were kept on ice all the time. To ensure the reproducibility of the measurements, we always use fluorescent calibration beads (Sphero Rainbow calibration beads 6 peaks; Catalog no. RCP-60-5; Spherotech) to adjust the instrumentation parameters.

### Live-cell confocal imaging of single mRNAs with blue light illumination

HeLa-ABC2 cell were plated on 3.5 cm glass-bottom FluoroDishes (FD35PDL, MPI) at a density of 3×10^4^ cells/dish one day in advance. Right before imaging, the medium was supplemented with 10mM HEPES, pH 7.4, and 1% penicillin/ streptomycin and placed on an Andor CSU-X1 spinning disk confocal with a Hamamatsu Orca-Fusion sCMOS camera, Nikon 60X NA1.49 TIRF objective, and Metamorph software. The blue light illumination was setup by inserting of Arduino controlled shutter (Daheng Optics GCI-7103) and blue filter (Omega 465/40DF) in the light path of bright field illumination. The brightfield illumination was adjusted to a blue light spot of 7 mm diameter centered at the bottom of the 35-mm plate, with light intensities of 25 or 100 μW/cm^2^. Multiple XY-positions within a 3 mm region were chosen, so every cell in the optical fields was always within the 7mm blue illumination circle, separated enough to avoid overlapping in excitation and photobleaching. At each XY-position, 20 z-slices with space of 0.5 μm was selected with 200 ms exposure per slice. A Prior piezo-z stage was used to ensure z precision. Each XY-position was imaged every 10 minutes for 1000 minutes with 37°C and 5% CO2. Excited by 561 nm laser, the laser power was selected to ensure the detection of single mRNA puncta while minimizing photobleaching over 100 3-D imaging loops. The emission filter 617/73 was used to collect tdTomato fluorescence. The AM light of 100 or 25 μW/cm^2^ or PWM with 200 minutes period, 25% on duty, and 100 μW/cm^2^ max intensity was set up and start right before the live-cell confocal imaging start. The period of 200 minutes was chosen over 400 minutes to enable multiple cycles of light inductions without significant photobleaching. It also significantly reduces noise over the AM (Figure 1-figure supplement 3B-C).

### Whole-genome sequence of HeLa-AB1 cell

The whole genome sequencing (50X) of HeLa-AB1 clone was performed by GENEWIZ (Suzhou, China). The reads were processed to call potential plasmid insertion sites by GENEWIZ (Suzhou, China). The reads were trimmed with cutadapt 1.9.1 to remove dual ends from reads with base quality cutoff of 20 and remove adapters. The trimmed reads were filtered with cutadapt 1.9.1 to discard reads containing more than 10% N bases and discard reads, which were shorter than 75 bp. The filtered reads were mapped to the reference genome (hg19) and plasmid insertion sequences using bwa 0.7.12-r1039. The alignments were sorted by leftmost coordinates using samtools 1.6. The duplicated reads in sorted alignments were marked with the MarkDuplicates function of picard 2.2.1. The break sites between the hg19 and plasmid were called from the alignments using VIFI 0.2.13. These sites were potential plasmid insertion sites in hg19 coordinates. PCR and Sanger sequencing were used to confirm the plasmid insertion sites. We confirmed 9 insertion sites for plasmid B1 and 5 for plasmid A1 (Figure 3-figure supplement 1B). The plasmid insertion sequences were inserted to the human reference genome (hg19) according to the validated insertion sites to build the single cloned cell line reference genome (HeLa-AB1) using a custom Python3 script. Every insertion site was updated after the front sites in the same chromosome had been inserted

### Chromatin immunoprecipitation-sequencing (ChIP-seq) and ATAC-seq sample preparation

HeLa-AB1 cell were plated in 10 cm dish at a density of 1.5×10^6^ cells/dish. The medium was supplemented with 1 μg/mL doxycycline on day −1 to induce GAVPO expression. On day 0, cells were illuminated by 25 μW/cm^2^ blue light for 24 hours to generate large mRuby distribution dispersion (expression heterogeneity). Cells were monodispersed and filtered through strainer to remove clumps. FACS Aria SORP was used to sort cells with high and low mRuby populations. Cells were kept on chilled water before and during sorting. Single cells were selected based on side and forward scattering properties. Manual gates were imposed on the mRuby fluorescence to collect approximately 20% of the lowest and highest mRuby-expression cells (Figure 3G). For each population, 5×10^6^ cells were collected. A fraction of the cells (∼2 ×10^5^) were cryopreserved with DMEM supplemented with 40% FBS (Gibco) and 10% DMSO (Catalog no. D8418; Sigma) for ATAC-seq analysis (GENEWITZ, Suzhou, China). The rest of the cells were washed twice in cold PBS buffer and cross-linked with 1% formaldehyde for 10 minutes at room temperature and then quenched by adding glycine (125 mM final concentration) for subsequent ChIP-seq experiments.

### ChIP-seq

The following ChIP-seq experiment is performed at IGENBOOK Biotech (Wuhan, China). Samples were lysed and chromatins were sonicated to obtain soluble sheared chromatin (average DNA length of 200–500 bp). Twenty microliters of chromatin were saved at –20°C for input DNA, and 100 μL chromatin was used for immunoprecipitation. The antibody used for H3K27me3 was ab4729 (Abcam). Ten micrograms of antibody was used in the immunoprecipitation reactions at 4 °C overnight. The next day, 30 μL of protein-A beads were added and the samples were further incubated for 3 h. The beads were then washed once with 20 mM Tris/HCL (pH 8.1), 50 mM NaCl, 2 mM EDTA, 1% Triton X-100, 0.1% SDS, twice with 10 mM Tris/HCL (pH 8.1), 250 mM LiCl, 1 mM EDTA, 1% NP-40, 1% deoxycholic acid, and twice with 1×TE buffer (10 mM Tris-Cl at pH 7.5, 1 mM EDTA). Bound material was then eluted from the beads in 300 μL of elution buffer (100 mM NaHCO3, 1% SDS), treated first with RNase A (final concentration 8 μg/mL) for 6 h at 65°C and then with proteinase K (final concentration 345 μg/mL) overnight at 45°C. Immunoprecipitated DNA was used to construct sequencing libraries following the protocol provided by the NEXTFLEX ChIP-Seq Library Prep Kit for Illumina® Sequencing (NOVA-514120, PerkinElmer Applied Genomics) and sequenced on Illumina Xten with PE150 method.

### ATAC-seq

The following ATAC-seq experiment is performed at GENWITZ (Suzhou, China). Resuspend cell pellet using 50 μL of tagmentation mix (12.5 μL 4X THS-seq TD buffer, 5 μL 0.1% Digitonin (Catalog no. G9441; Promega), 5 μL TTE Mix V50 (Catalog no. TD501; Vazyme), 27.5μl H2O). The tagmentation reaction was performed on a thermomixer (Eppendorf 5384000039) at 800 rpm, 37 °C, 30 min. The tagmented DNA was purified following Qiagen MinElute PCR purification kit protocols. Library was constructed using 10 μL purified tagmented DNA, 2.5 μL S5XX and 2.5 μL N7XX indexing primers, 25 μL of NEBNext 2X PCR MasterMix and 10 μL H2O, amplified with 72 °C 5 min, 98 °C 1 min, [98 °C 10 s, 63 °C 30 s, 72 °C 20 s] × 12. The product was purified using SPRI beads method (Beckman Coulter). Sequencing was performed on an Illumina HiSeq with PE150 method.

### Single cell ATAC-seq

The period of 600 minutes for PWM light induction was chosen as it reduces noise and better fitted for covering chromatin dynamics studies from 0 to 1 day (Figure 4A). Briefly, 50,000 cells were centrifuged down at 500×g, 4 °C, 5 min. Cell pellets were resuspended in 50 μL tagmentation mix (33 mM Tris-acetate, pH 7.8, 66 mM potassium acetate, 10 mM magnesium acetate, 16% dimethylformamide (DMF), 0.01% digitonin and 5 μL of Tn5 from the Nextera kit from Illumina, Cat. No. FC-121-1030). The tagmentation reaction was performed on a thermomixer at 800 rpm, 37 °C, 30 min. The reaction was then stopped by adding equal volume (50 μL) of tagmentation stop buffer (10 mM Tris-HCl, pH 8.0, 20 mM EDTA, pH 8.0) and left on ice for 10 min. A volume of 200 μL 1X DPBS with 0.5% BSA was added and the nuclei suspension was transferred to a FACS tube. Tagmented single nuclei were sorted based on GFP signal (nuclear-localized tetR-GFP-nuc) using BD FACSAria SORP into one prepared 384-well plate containing 4 μL lysis buffers (50 mM Tris, pH 8.0, 50 mM NaCl, 20 ug/ml Proteinase K, 0.2% SDS, 10 μM S5xx/N7xx Nextera index primer mix (5 µM each)).

After sorting, the plate was briefly centrifuged and incubated at 65 °C for 30 min. Then, 4 μL 10% tween-20, 2 μL H2O and 10 μL NEBNext® High-Fidelity 2 ×PCR Master Mix was added to each well sequentially. Libraries were amplified with 72 °C 5 min, 98 °C 5 min, [98 °C 10 s, 63 °C 30 s, 72 °C 20 s] × 18. Finally, all reactions were pooled together and purified with a PCR minElute purification column (Qiagen). Libraries were sequenced at HaploX Biotech (Shenzhen, China) with a HiseqX machine with 40G per one 384-well plate.

### Single cell RNA-seq Library preparation

HeLa-AB1 and F9-AB2 cells were seeded to in ibidi 24-well μ-plate at 30,000 cells per well one day prior to experiments. At day 0, two wells (A2, A3, control) and one well of HeLa-AB1 cells (C2, inhibitors) were replenished with regular media and regular media containing 0.333 μM LMK-235 and 1 μM A-485, respectively, and illuminated with blue lights for two days at 21 and 9 μW/cm^2^, respectively. At the same time, two wells (A1, B1, control) and one well (C1, inhibitors) of F9-AB2 cells with regular media and regular media containing 1 μM LMK-235 and 1 μM A-485, respectively, and continue culture at dark for 2 days. F9-AB2 cells (three samples named A1, B1 and C1 respectively) and HeLa-AB1 cells (three samples named A2, A3 and C2, respectively) in ibidi 24-well μ-plate were washed with 500 μL DMEM for three times, following incubated in 125 μL TrypLE Express Enzyme at 37 °C for 5 minutes. This enzyme was deactivated by 250 μL DMEM+10%FBS and the cells were transferred into 1.5 mL centrifuge tubes and centrifuged down at 200×g, 4 °C for 5 min. Cell pellets were washed with 1 mL PBS with 0.1% w/v BSA. Discard the PBS as much as possible and resuspend the cell pellets with an additional 100 μL PBS containing 0.1% BSA. The cells were filtered through the filter cap of flow tube (Catalog no. 352235; BD Falcon). Aliquots of 10 μL filtered cells were mixed with equal volumes of trypan blue solution for cell counting with the Countstar analyzer (RuiYu Biotech, Shanghai, China). Finally, the human cells (HeLa-AB1 A2, A3, and B2) and mouse cells (F9-AB2, A1, B1 and C1) were mixed at 1:1 ratio to form three samples (A1/A2, B1/A3, C1/C2). Subsequently, according to the 10× Genomics user guide, scRNA-seq libraries were constructed in three steps including GEM Generation & Barcoding, Post GEM-RT Cleanup & cDNA Amplification and 3’ Gene Expression Library Construction. In the first step, in order to obtain a final recovery of 10,000 cells/sample, 16,500 cells/sample were loaded into the 10× machine (Chromium Next GEM Single Cell 3’ Library & Gel Bead Kit v3.1). In the third step, after the scRNA-seq libraries constructed, each library is split into two sets. One set was sequenced on HiseqX system at HaploX Biotech, while the other was sequenced on MGI2000 platform at BGI.

### Flow cytometry data analysis

The flow cytometry data were gated for single-cell and exported as csv files using FlowJo X (Ashland, US). A custom MATLAB (Mathworks, Natick, US) script loads the csv files for quantitative analysis, including mean, CV, histogram, etc. The relevant fluorescent channel was normalized to the mean value of the fifth peak of the Rainbow calibration beads. There is no detectable spillover between GFP (FITC-A), mRuby (ECD-A), and BFP (PB450A) channel, so compensation was not needed.

### ChIP-seq and ATAC-seq data analysis

Reads were preprocessed using the fastp 0.20.0 with the default setting and correct low-quality mismatched base pairs in overlapped regions of paired-end reads. The preprocessed reads were mapped to the single cloned cell line reference genome (HeLa-AB1) using bowite2 2.3.5.1. The alignments were preprocessed with samtools 1.9 and then deduplicated using the MarkDuplicates function of picard 2.22.2. The deduplicated alignments were filtered using samtools 1.9 with SAM flag (-F 1804 -f 2) to select reads mapped in proper pair. These reads were filtered using custom bash code to remove the reads which were mapped to mitochondria. For population sequencing data, all samples filtered reads were processed using the multiBamSummary function of the deeptools 3.4.2 to calculate the scale factor for each sample. The scale factor and filtered reads for each sample were processed using the bamCoverage function of deeptools 3.4.2 to calculate the normalized coverage track. For single-cell sequencing data, the filtered reads were processed using the bamCoverage function of deeptools 3.4.2 to calculate the coverage track. The coverage tracks were processed using the custom python3 scripts to calculate the coverage around the TSS sites.

### scRNA-seq data analysis

The human and mouse mixed reference file was from the 10x genomics website (https://cf.10xgenomics.com/supp/cell-exp/refdata-cellranger-hg19-and-mm10-3.0.0.tar.gz). The coding region of vector CEG was added to the reference file and make a new reference with the function mkref of the cellranger 3.1.0. The human and mouse mixed reads were processed by cellranger 3.1.0 to obtain the UMI count matrix. The matrix files were processed by R package scater 1.16.2 using the function quickPerCellQC with the default setting to filter low quality cells. The cells were identified as human cells as if the total UMI counts mapped to mouse genome were less than the 0.01 times of the total UMI counts mapped to human genome. The cells were identified as mouse cells as if the total UMI counts mapped to human genome were less than the 0.018 times of the total UMI counts mapped to mouse genome. The UMI counts were normalized by the library size. All the following computations in this section were performed using MATLAB R2018b and associated toolboxes. The genes with mean counts under 1 CPM (counts per million) in any sample were neglected. Coefficients of variance (CV) and means for the remain genes were computed for all three samples. The CV vs mean dot scatter plot was fitted to a power function (“power1”) using the fit function. A cutoff line for identifying genes with CV outliers was defines as 3 σ above the fitted curve.

The σ for each mean level is computed as the standard deviation of the CVs for the genes with similar mean. The genes above this cutoff line (mean > 5 CPM and CV > 1.0), in any of the three samples, were filtered out for further analysis. The normalized count matrices from two control samples were pooled together as the control. If the CV of a gene is smaller than 2 in the control, it is grouped in the “low CV” group, otherwise it is grouped in the “high CV” group. Transcription factor (p300, CBP and p65) target gene sets were collected from the ChIP-Altas potential target genes with the option of ±1kb distance from TSS. Genome assembly mm10 was selected as mouse reference in ChIP-Altas. Genome assembly hg19 was selected as human reference in ChIP-Altas. The UMI counts of filtered genes were processed by the R package saver 1.1.2 (Huang et al., 2018) to get a smooth distribution (Figure 6G).

### Imaging processing of single mRNA imaging

MCP-tdTomato-nuc fluorescence is mostly localized inside the nucleus, which enables us to segment, track individual nucleus, detect mRNA in single channel (Figure 5A). Each optical field was captured to 4D XYZT image with 1300 pixels × 1900 pixels × 20 slices × 100 timepoints. Mapping with the camera pixel size of 6.5 μm, 60X NA1.49 TIRF objective, 0.5 μm per z-slice, and imaging frequency of every 10 minutes, each 4D images recorded 140×205×9.5 μm^3^ over 1000 minutes. The cells move significantly over 1000 minutes, and most of the nuclei went out of the imaging boundary at one time or another. To find all nuclei stay entirely inside the boundary, we performed nuclear segmentation and tracking before using Imaris 9.4 to identify and count mRNAs. The first step is generating a z-axis maximum projection. A medium filter (window size=5) and Laplacian of Gaussian filter (sigma=40) were applied sequentially to projected XYT images, and a threshold was chosen to binarize the images and identify nuclei’s centroid. Connected components were searched in these binary images over time to track the nuclei remained inbound. In addition, for each z-projection image, using the identified nuclei centroid as seeds, a watershed algorithm is applied to find the “basin” occupied by each cell at every time points. For all the tracked nuclei, the entire cell region was cropped from the original XYZT images based its “basin” at each time points, re-centered to generated in a 512×512×20×100 box. The nucleus is finally segmented using Otsu’s algorithm in the cropped image. These XYZT images are normalized to the mean nuclear fluorescent intensity to compensate for MCP-tdTomato-nuc expression heterogeneity and possible photobleaching, as Imaris doesn’t incorporate adaptive thresholding. The single mRNAs were identified using Imaris’ spot model with an estimated xy diameter of 0.54 μm and z diameter of 1.5 μm. All the codes were written in Julia 1.4.1 using the JuliaImages v0.20 package.

### Statistical Analyses

Data are described by the sample mean and standard deviation (SD). All statistical analyses were performed in MATLAB 2018b statistic toolbox. Sample size, details of statistical tests are provided in the corresponding figure legends.

## Acknowledgments

This work was financially supported by the National Key R&D plan project (2019YFA0906002), National Natural Science Foundation of China projects (31971181, 31770928), Project KQTD2016053117035204 Supported by Shenzhen Peacock Plan, Science and Technology Planning Project of Guangdong Province, China (2016A050503010). We appreciate professors Qing Nie, Chunhui Hou, Wei Chen, and Liang Fang for stimulating discussions and suggestions, Mr. Wang Yuhao, Yi Zhang and Jiashun Xiao and other members of the Huang group for technical assistant, Miss. Yifan Zhu for help with illustration. We appreciate professors Yi Yang, Jun Chu, and Jiandong Huang, Jianbo Yue for providing plasmids for lightON, mRuby, tetR negative feedback plasmids, and mouse ES D3 cell. We would like to acknowledge the technical support of Dr. Wenjie Wei from SUSTech Research Core Facility.

## Funding

Funder Grant reference number Author

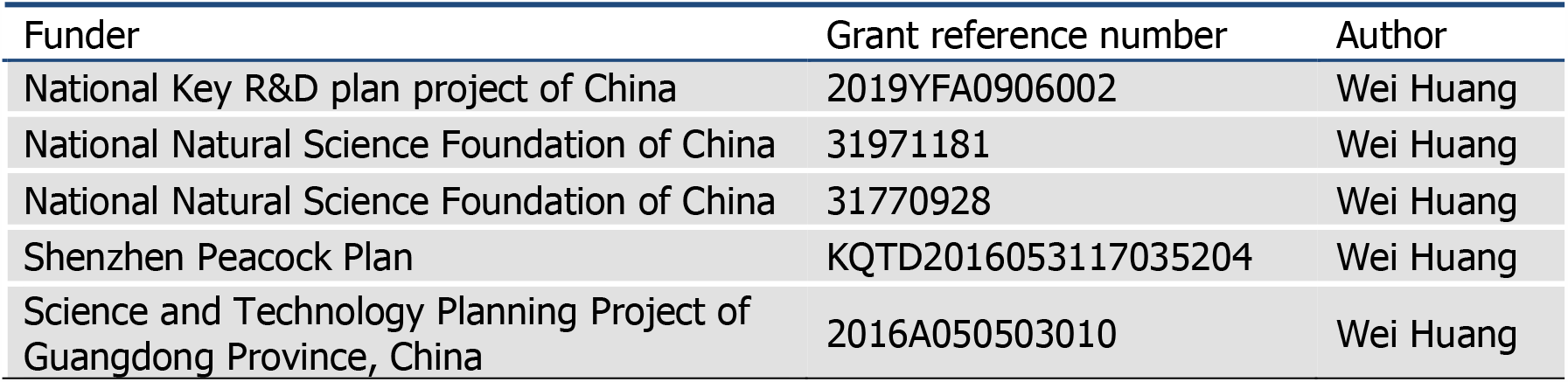

## Author contributions

Conceptualization: W.H.; molecular and cell biology experiments: D.T., X.L., H.H., S.G., W.F.; light modulation system: Y.M., F.J., H.H., and W.H.; microscopic imaging and image analysis: Y.M., Y.W., D.T. and W.H.; epigenetic experiments: D.T., W.X. and X.C.; epigenetic data analysis: R.C., and X.C., other data analysis, and modeling: R.C., H.H., and W.H.; writing: W.H.; editing: D.T., R.C., Y.M., W.X., X.C. and W.H.

## Additional files Supplementary files

### Data and Code Availability

Sequencing data has been deposited in the European Nucleotide Archive under the accession-number PRJEB40269 and PRJEB41743. Scripts for sequencing data analysis are available at GitHub (https://github.com/QBioLab/sequence-data-analysis-for-noise-control. Live cell mRNA image processing codes are available at GitHub (https://github.com/QBioLab/CountmRNA.jl). Simulation and analysis codes for the ODE model is available at GitHub (https://github.com/QBioLab/noise-simulation).

## Supplementary Figures

**Figure 1-figure supplement 1.**
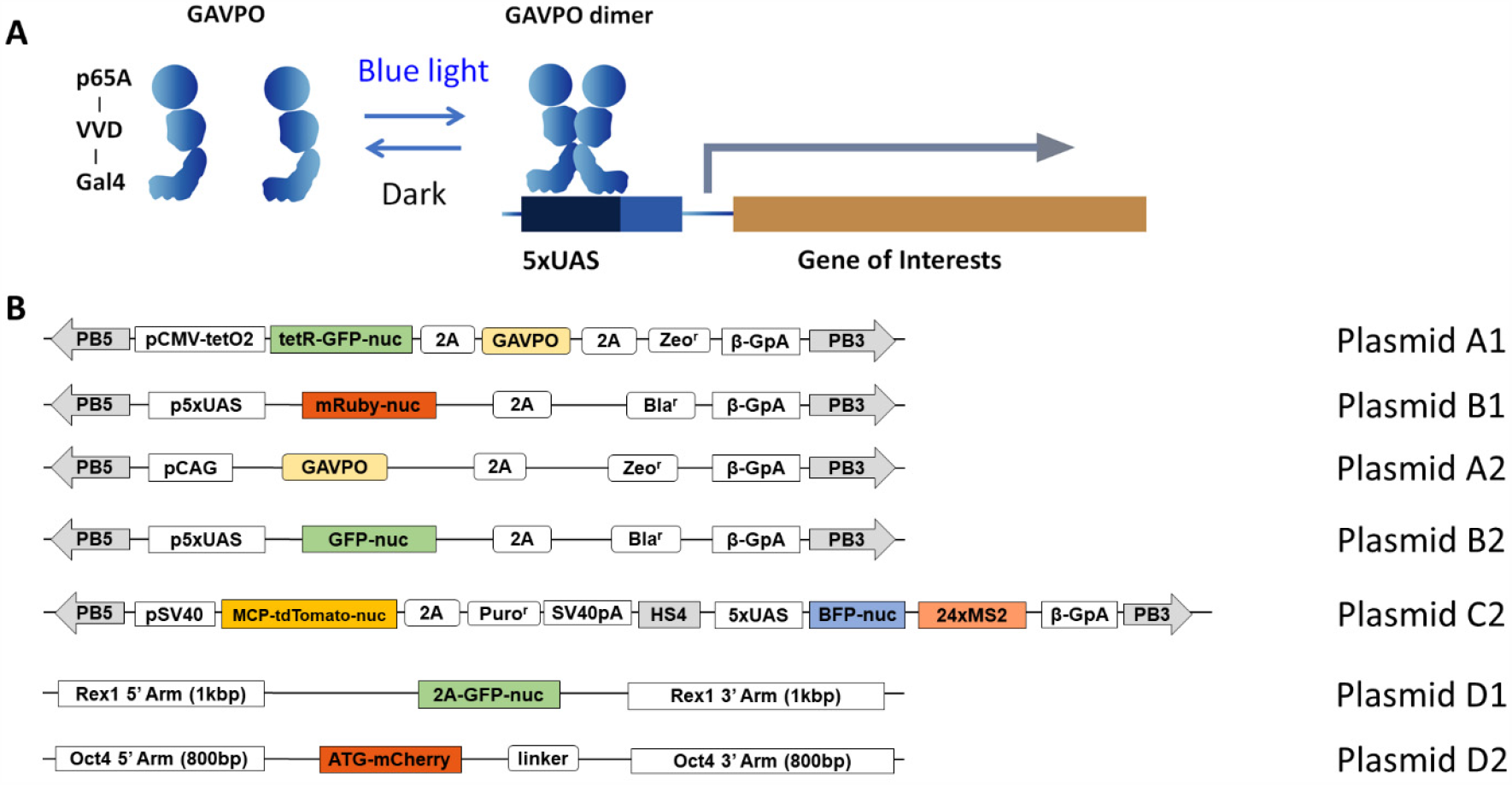
Plasmids designs. (**A**) schematic view of the lightON expression system. (**B**) plasmids constructed for lightON expression circuits. PB5 and PB3 are the two terminal repeats for PiggyBac transposon. pCMV-tetO2 is the engineered pCMV promoter with two tetO sequences sandwiched the TATA box. tetR-GFP-nuc is the fusion of tet repressor, EGFP, and 3 repeats of SV40 nuclear localization sequences. pCAG is the CAG promoter, pSV40 is the SV40 promoter. All the other symbols are described in detail in the method section.

**Figure 1-figure supplement 2.**
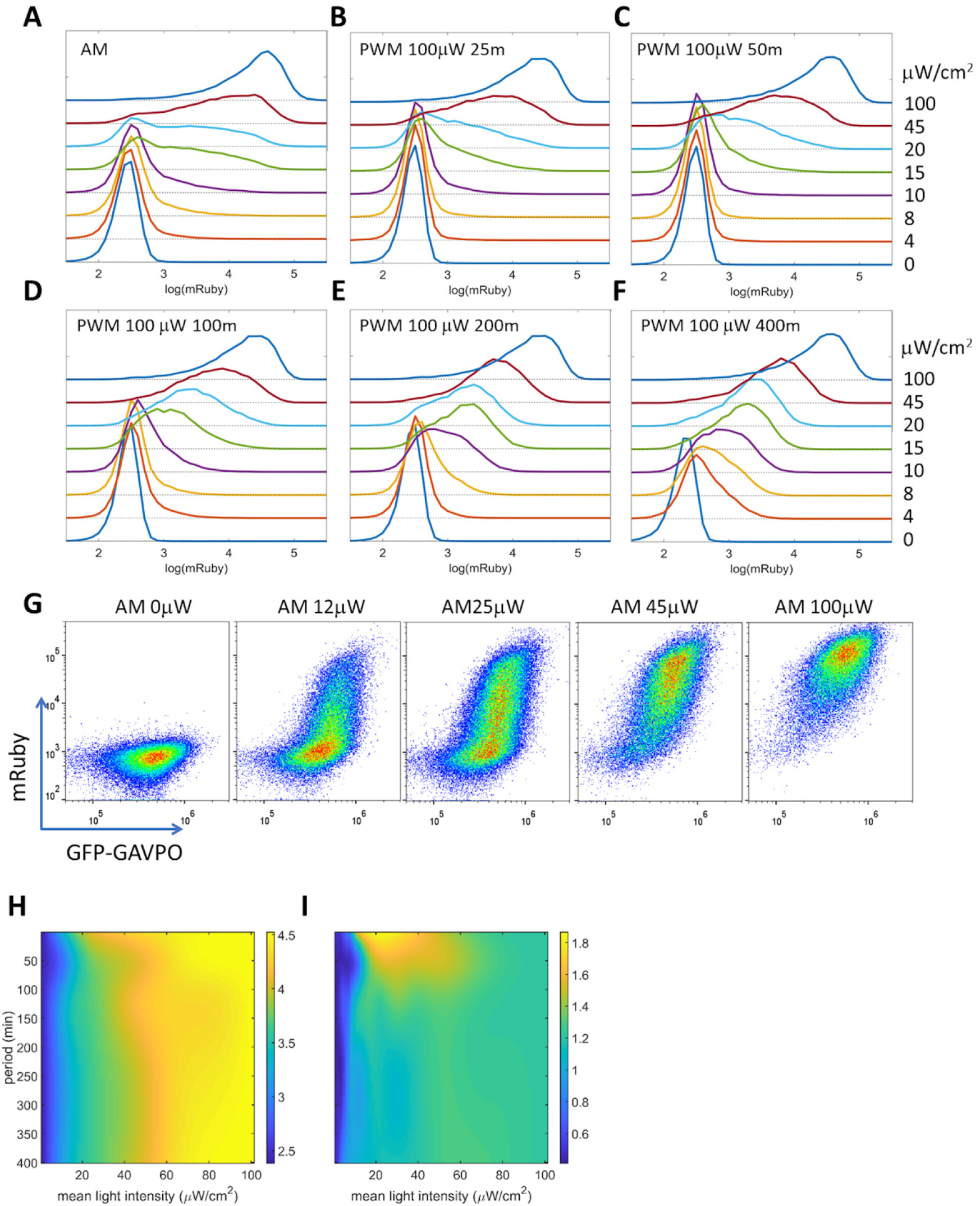
PWM modulation of distribution dispersion in HeLa-AB1 cell. **(A-F)** Histograms of mRuby expression of HeLa-AB1 cells under AM light (A), PWM light with periods of 25 m (B), 50 m (C), 100 m (D), 200 m (E) and 400 m (F). The light intensities (AM) or duty (ON fraction) are labeled to the right of t the plots. (**G**) The mRuby-GFP-GAVPO density plots for HeLa cell populations under AM light with 0, 12, 25 μW/cm^2^, 45 μW/cm^2^ and 100 μW/cm^2^. Surface response plots of (H) mean mRuby over the period and mean light intensity of PWM induction, and (I) mRuby distribution spreading (ratio of mRuby intensities of 90 percentile to 10 percentile) over the period and mean light intensity of PWM induction. Each sample contains 10,000-50,000 cells.

**Figure 1-figure supplement 3.**
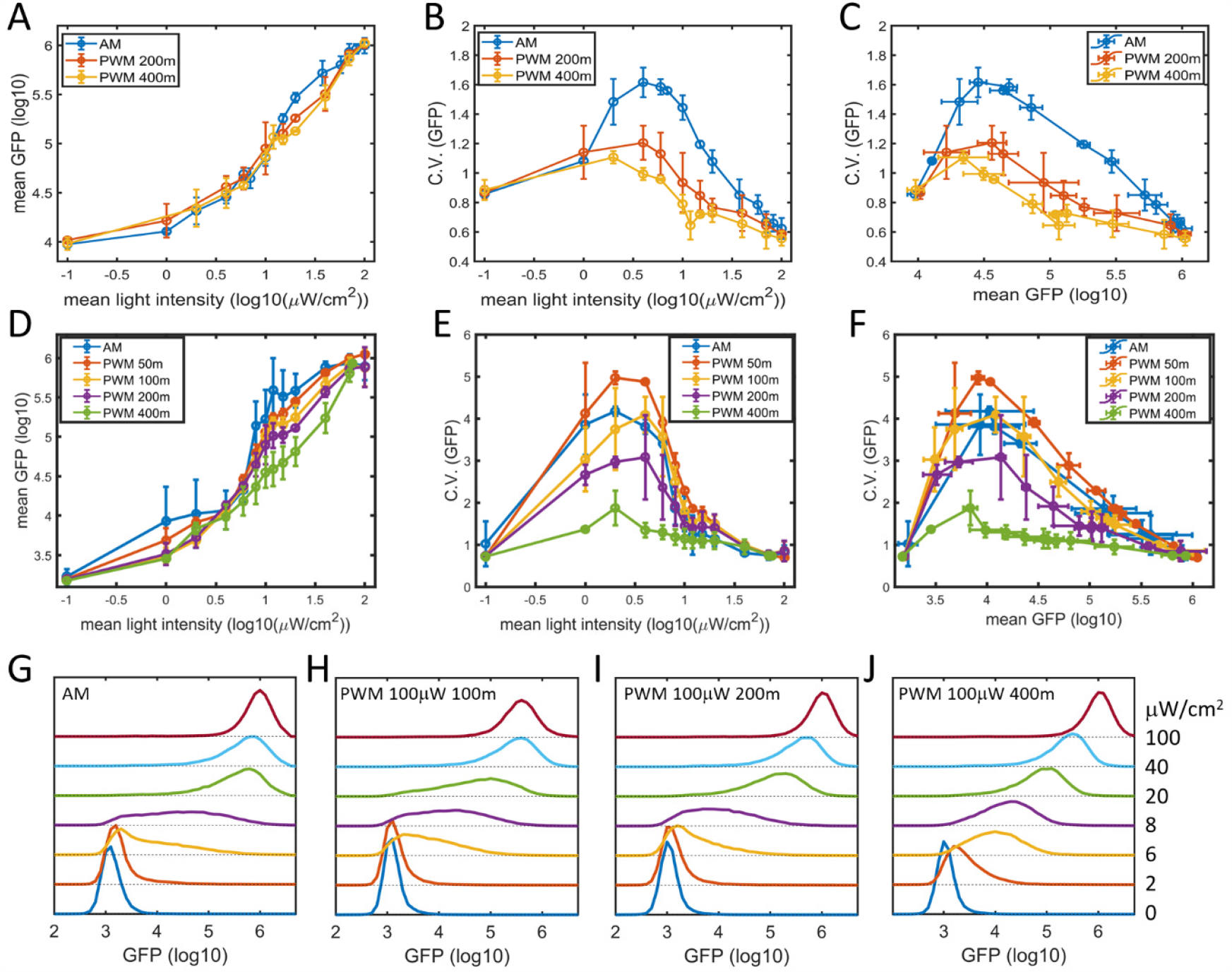
Noise modulations in HeLa-AB2 and F9-AB2 cells. (**A**) Mean GFP versus mean light intensities, (**B**) CVs of GFP versus mean light intensities, (**C**) CVs of GFP versus mean GFP light intensities for HeLa-AB2 clone under AM light (blue), PWM light with periods of 200 m (red), and 400 m (yellow). (**D**) Mean GFP versus mean light intensities, (**E**) CVs of GFP versus mean light intensities, (**F**) CVs of GFP versus mean GFP light intensities for F9-AB2 clone under AM light (blue), PWM light with periods of 50 m (red), 100 m (yellow), 200 m (purple) and 400 m (green). Error bars represent standard deviations for at least two independent experiments. (**G-J**)The histograms of GFP for F9-AB2 cells were induced with AM lights (G) and PWM lights with periods of 100 m (H), 200 m (I), and 400 m (J). The mean light intensities are to the right. Each sample contains 10,000-50,000 cells. The error bars represent standard deviations from 2-3 independent experiments.

**Figure 2-figure supplement 1.**
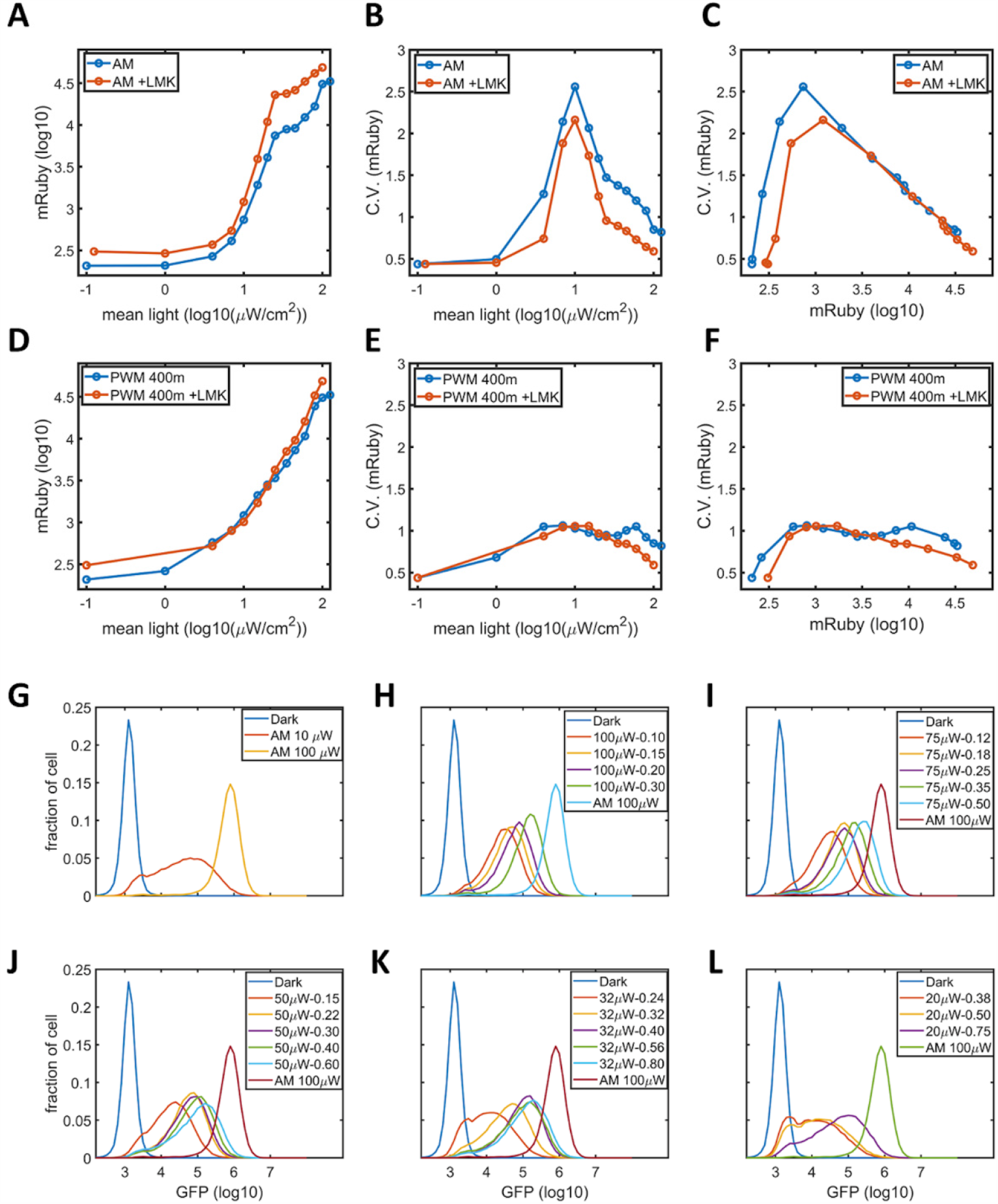
Evidences for epigenetic bistability. (**A**) The mean mRuby against mean light intensities, (**B**) CVs of mRuby against mean light intensities, (**C**) CVs of mRuby against mean mRuby for HeLa-AB1 cells under AM light, without LMK-235 (blue line) and with 0.333 μM LMK-235 (red line) for two days. (**D**) The mean mRuby against mean light intensities, (**E**) CVs of mRuby against mean light intensities, (**F**) CVs of mRuby against mean mRuby for HeLa-AB1 cells under PWM light with a period of 400m, without LMK-235 (blue line) and with 0.333 μM LMK-235 (red line) for two days. (**G-L**) The histograms of GFP for F9-AB2 cell with AM light (G), PWM light with period of 400 m and maximum light intensities of 100 μW/cm^2^ (H), 75 μW/cm^2^ (I), 50 μW/cm^2^ (J), 32 μW/cm^2^ (K), and 20 μW/cm^2^ (L). The histograms for Dark and AM with 100 μW/cm^2^ are used as references. Each sample contains 10,000-50,000 cells.

**Figure 2-figure supplement 2.**
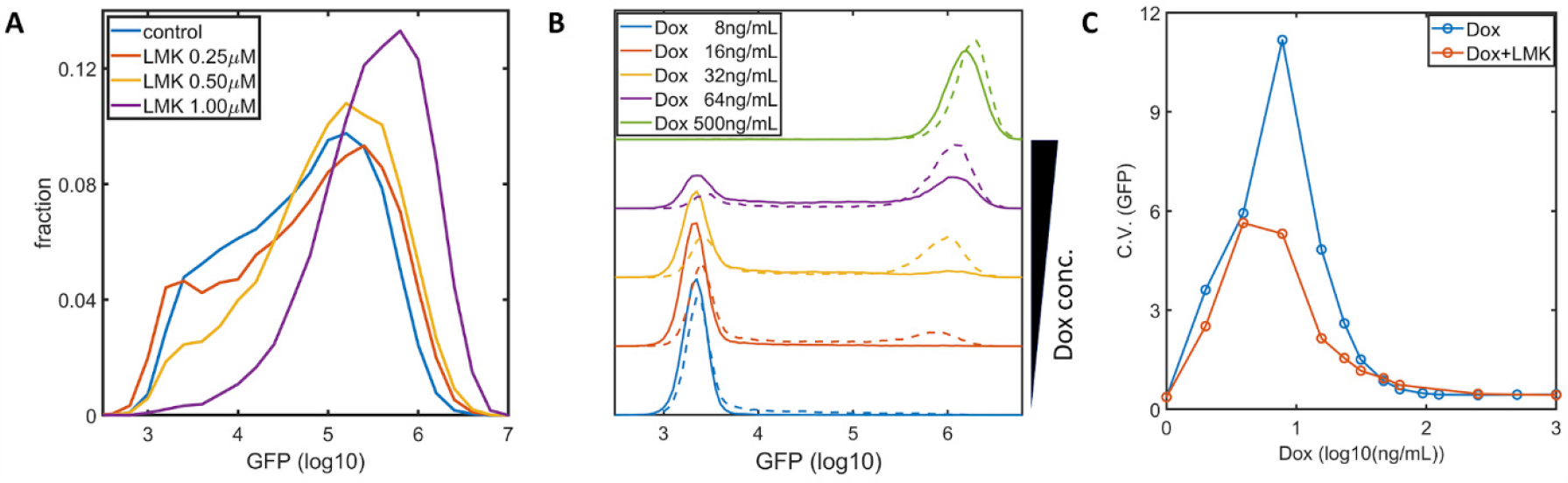
Inhibition of HDAC4/5 on F9-AB2 and HeLa-Tet-On cells. (**A**) The histograms of GFP expression with F9-AB2 cells with AM light (10 μW/cm^2^) alone (blue), AM light with 0.25 μM (red), 0.5 μM (yellow), and 1 μM (purple) LMK-235. (**B**) Histograms of GFP for HeLa-Tet-On cells (stable transfected with Tet-On-GFP expression circuit(Lu and Huang, 2014)) with doxycycline alone (solid lines), and doxycycline plus 0.333 μM LMK-235 (dashed line). The doxycycline concentrations are labeled inside the plots. (**C**) CVs of GFP for HeLa-Tet-On cells with doxycycline alone (blue circles), and Doxycycline plus 0.333 μM LMK-235 (red circles). Each sample contains 10,000-50,000 cells.

**Figure 3-figure supplement 1.**
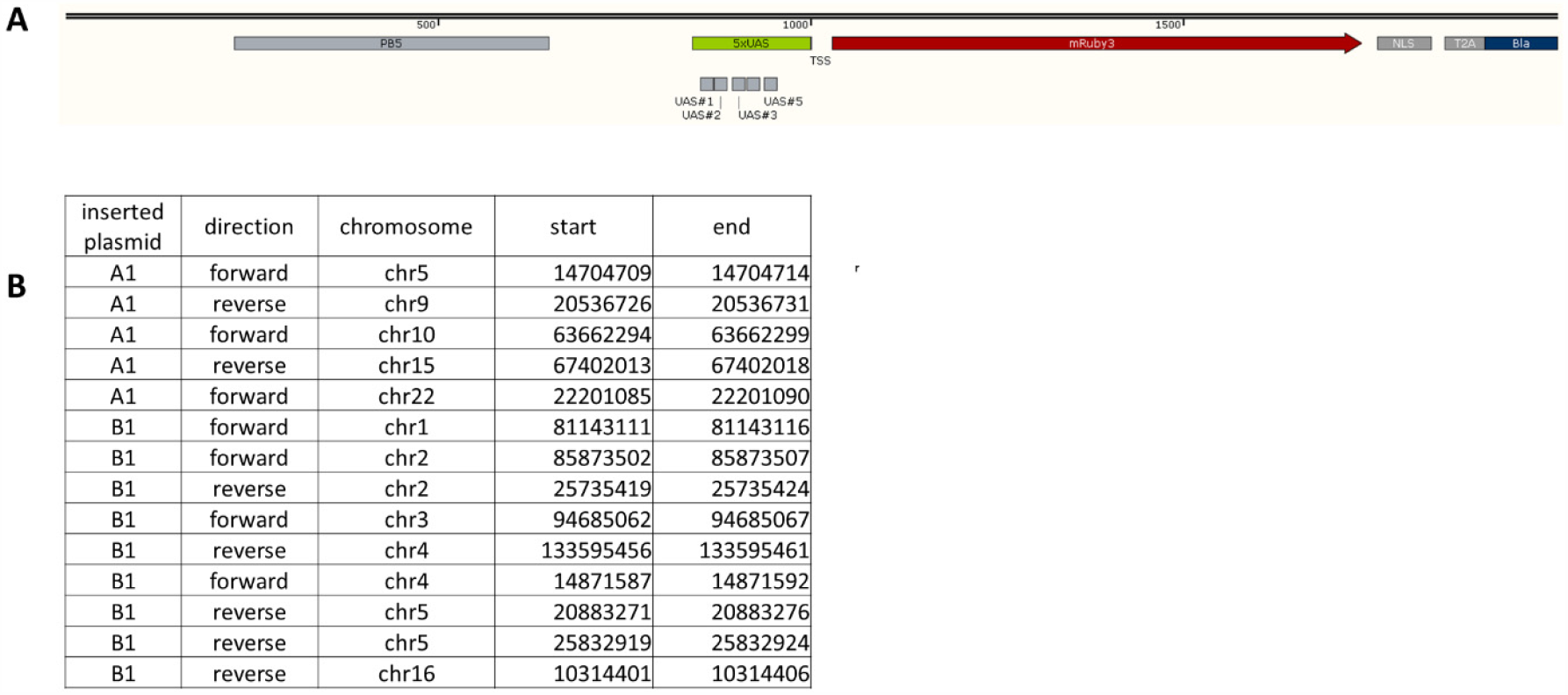
Insertion sites of the lightON cassettes. (**A**) Part of the inserted sections of the lightON cassette (plasmid B1), with TSS aligned at 1000 bp. The inserted sequences are between PB5 and PB3 (not shown in the diagram). (**B**) The insertion sites for plasmids A1 and B1 in HeLa-Ab1 cell, based on 50X whole genome sequence and PCR/Sanger sequencing confirmation.

**Figure 5-figure supplement 1.**
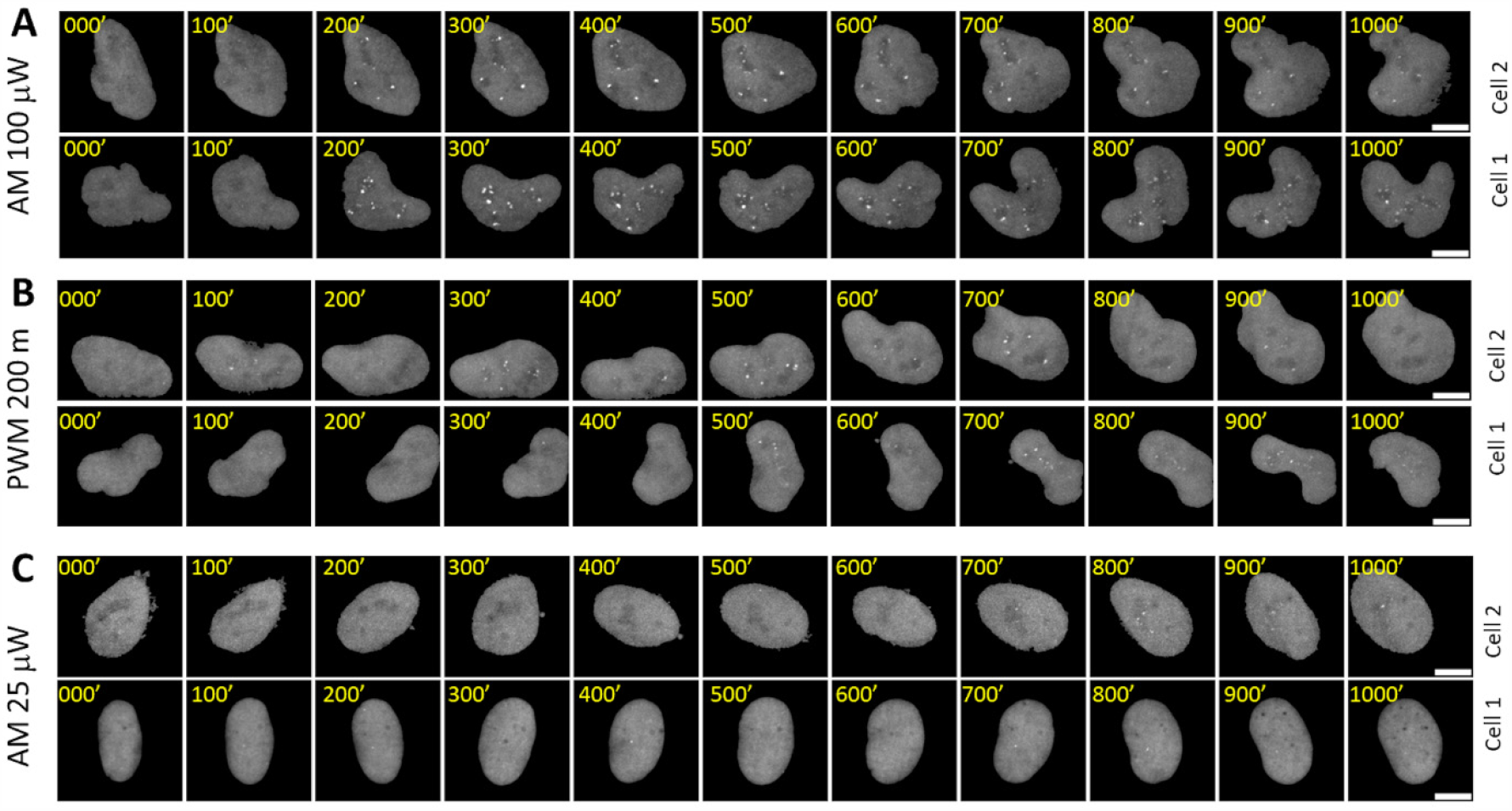
Time lapse images of representative nuclei. (**A-C**) The z-projection images of two representative cell nuclei over 1000 minutes each are displayed for AM with 100 μW/cm^2^(A), PWM with 100 μW/cm^2^ and a period of 200 m (B), and AM with 25 μW/cm^2^(C). The nucleus was segmented, tracked, isolated, and re-centered as described in the method section. The scale bars represent 10 μm.

**Figure 6-figure supplement 1.**
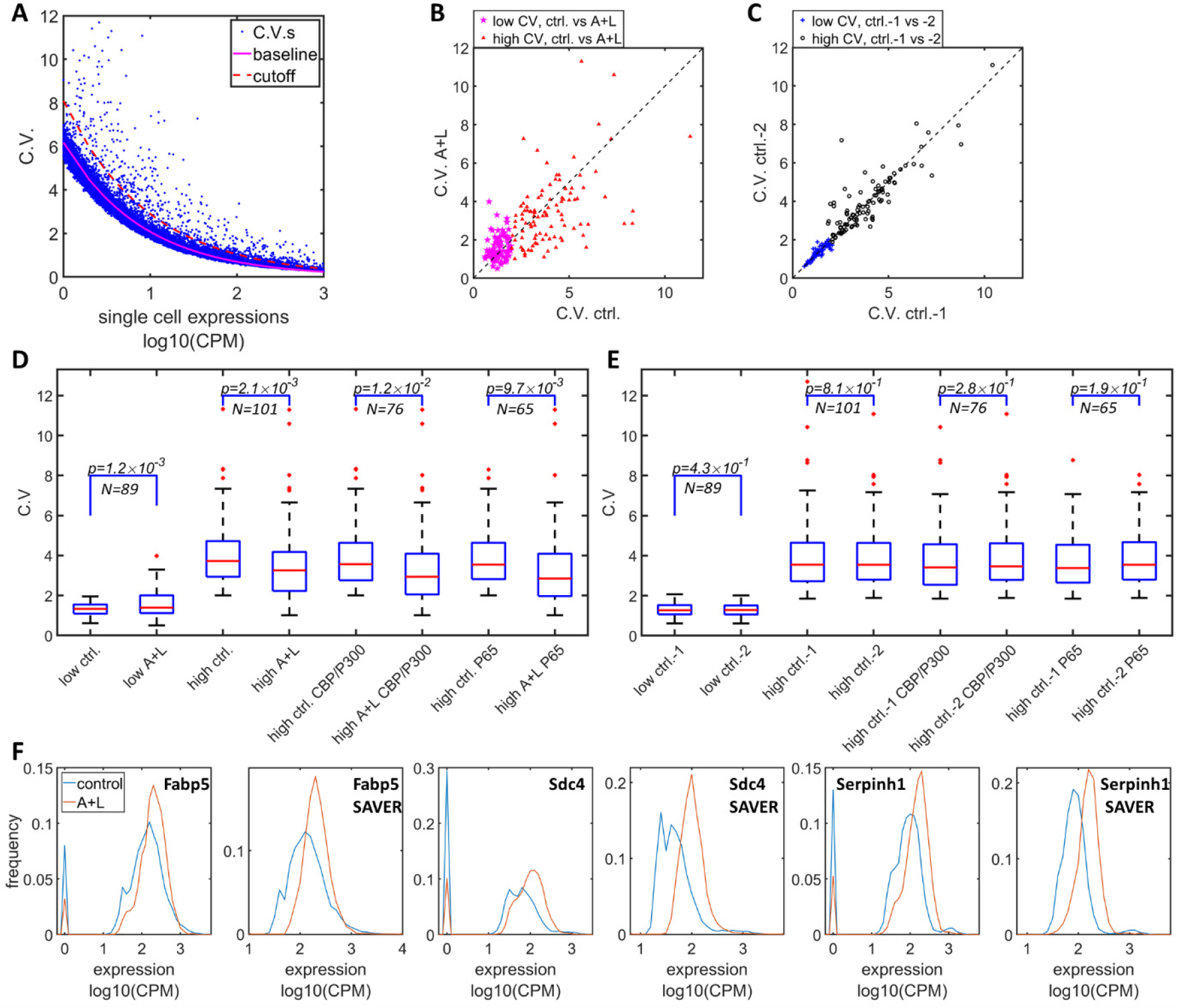
Simultaneously inhibition of CBP/p300 and HDAC4/5 reduces heterogeneity in endogenous gene expressions in F9 cell. (**A**) The CV vs mean expressions plot with scRNA-seq data from F9 with or without 1 μM LMK and 1 μM A-485. Purple solid line represents the fitted power function of all the genes, with exponent of −0.47. Red dashed line represents the 3-sigma cut off for noisy outlier genes. (**B**) The genes with CV vs gene expression above the cutoff (red line in C), and mean reads >5CPM were selected, and plotted as CV between F9 cell with A485+LMK-235 and control condition. Purple pentastars and red triangles represent the genes with low (< 2.0) and high CV (≥2.0), respectively. (**C**) Blue asterisks and black circles represent the CV of the same gene sets between two biological replicates of control cells. (**D**) Statistical analysis of the CV of genes between A485+LMK-235 and control, with low CV, high CV, high CV and positive for CBP/p300 or p65 ChIP. The boxes show the lower and upper quartiles, the whiskers show the minimum and maximal values excluding outliers; the line inside the box indicates the median; outliers (red dots) were calculated as values greater or lower than 1.5 times the interquartile range. The p values calculated using paired student *t*-test is shown in the figure. (**E**) Similar statistical analysis of CV between two biological replicates of control conditions for the same sets of genes. (**F**) Histograms of expressions for control (blue) and A485+LMK-235 group (red) of genes Fabp5, Sdc4 and Serphinh1, calculated from normalized data without or with SAVER denoising algorithm. The peaks at “0” in the histograms represent cells with zero reads.

